# A novel ILK/STAT3 pathway controls plasticity in a neural stem cell model of glioblastoma

**DOI:** 10.1101/2023.07.19.549653

**Authors:** Alexander E. P. Loftus, Marianna S. Romano, Anh Nguyen Phuong, Morwenna T. Muir, John C. Dawson, Lidia Avalle, Adam T. Douglas, Richard L. Mort, Adam Byron, Neil O. Carragher, Steven M. Pollard, Valerie G. Brunton, Margaret C. Frame

## Abstract

Glioblastomas (GBM) are driven by malignant neural stem-like cells that display extensive heterogeneity and phenotypic plasticity, which drives tumour progression and therapeutic resistance. Here we show that the nodal extracellular matrix-cell adhesion protein integrin-linked kinase (ILK; a pseudokinase), is a key determinant of phenotypic plasticity and the mesenchymal-like, invasive cell state in mouse GBM stem cells. We found that a novel ILK-STAT3 signalling pathway is required for plasticity that enables the transition of GBM stem cells to an astrocyte-like state both *in vitro* and *in vivo*. GBM cells genetically depleted of ILK become predominantly stabilised in a transcriptionally-defined progenitor-like state that is characterised by lack of response to differentiation cues and constitutive proliferation. Loss of ILK or interference with STAT3 impairs differentiation potential, reducing phenotypic plasticity of tumour cell populations; additionally, ILK loss causes a mesenchymal- to epithelial-like morphological transition and suppression of malignancy-associated features. Our work defines ILK as a central regulator of multiple GBM phenotypes including phenotypic plasticity and mesenchymal state.

## Introduction

Glioblastoma (GBM) is the most common and aggressive malignant primary brain tumour in adults^1^, accounting for more deaths in those under 40 than any other tumour type^2^. Patient survival has not improved significantly for decades^3^; GBM is essentially incurable with median survival of ~15 months from diagnosis^4^. Although increasing knowledge of genetic and epigenetic changes has revealed the complex molecular etiology of GBM and has helped refinement of tumour sub-typing, it has not yielded tractable precision medicine hypotheses to improve clinical outcomes. No single actionable driver has been uncovered, as the key GBM signaling pathways can be disrupted at many levels. There is therefore an urgent need to better understand the biology of GBM, including the nature of pathways that regulate events during transformation including plasticity between distinct cell states, and formulate tractable hypotheses regarding vulnerabilities within or across sub-types^2^.

GBM is largely driven by cells with molecular and phenotypic hallmarks of neural stem cells (NSCs) that exploit developmental pathways to fuel tumour growth, recurrence, and therapeutic resistance^5, 6^. Recent data suggests that GBM stem cells hijack a core NSC injury response transcriptional program, and that adhesion genes such as β1 integrin and its downstream effector integrin-linked kinase (ILK), a pseudokinase^7^, may represent critical acquired vulnerabilities^6^. Indeed, abnormal integrin signalling contributes to the malignant features of GBM cells, including invasion, angiogenesis, extracellular matrix (ECM) degradation, immune modulation, and the maintenance of stem cells^8, 9^. Furthermore, ECM components, like laminin and fibronectin, and their cognate receptors, are important in GBM stem cell maintenance, proliferation and migration^10–12^. GBM cells adopt a mesenchymal-like mode of migration, relying on integrin-linked adhesion proteins to act as ‘molecular clutches’ that control force needed for movement through their environment^13^. ILK is a central player in the ‘consensus integrin adhesome’, derived by integrating integrin adhesion complex proteomes from diverse cell types^14^, which consists of 68 proteins present in each of the integrin-protein complex datasets examined. Four focal adhesion nexuses including a nexus centred around ILK and its binding partners PINCH-1 and ɑ-parvin were identified, the components of which link integrin complexes to actin and multiple intracellular signalling pathways, and may represent critical vulnerabilities in cancer^6^.

Here, we made use of our recently-described genetically-transformed NSC model of GBM^15^, featuring key driver mutations which initiates tumours and recapitulates the aggressive features of the disease^16–21^. We uncovered an unexpected role for ILK in regulating GBM stem cell plasticity *in vitro* and *in vivo*. Transcriptomic profiling of cells depleted of ILK revealed reduction of gene expression signatures that regulate differentiation and plasticity and a gene expression profile consistent with ILK-deficient GBM stem cells being predominantly retained in a progenitor stem cell state. Consistently, ILK depletion resulted in an inability of GBM stem cells to undergo differentiation into astrocytes (which we modelled using both BMP-4 and serum treatment^15, 22, 23^) and prevented exit from the cell cycle. We further show that ILK is required to induce phosphorylation of STAT3, and inhibiting this pathway genetically or pharmacologically impaired the BMP-4-induced cytostatic response. ILK-deficient GBM stem cells could grow when transplanted into the mouse brain, but the resulting tumours had distinctive properties characterised by suppressed astrocytic differentiation and STAT3 signalling, and reduced *in vivo* motility. Consistently, ILK caused marked changes in expression of genes associated with astrocyte-like and proliferative progenitor cell states *in vivo*. Finally, patient tumours enriched for the proliferative progenitor cell state featured reduced expression of ILK protein and its key binding partners. Our findings therefore show that ILK, via a previously unknown ILK-STAT3 signalling pathway, promotes the diverse developmental states seen in GBM and its loss abolishes signalling required to maintain GBM stem cell transcriptional plasticity. We discuss how trapping GBM cells in a restricted progenitor-like cell state by inhibiting the novel ILK-STAT3 pathway may be exploited to suppress cell state transitions that are linked to malignant behaviour.

## Results

### ILK controls morphology, transcriptional programming and astrocyte state transition in GBM stem cells

GBM arises from cells with stem-like characteristics that transition to various neurodevelopmental states^5, 6, 24^. To explore the role of ILK in GBM, we utilised a recently-described transformed NSC-derived GBM cell model^15^ featuring CRISPR/Cas9-mediated deletion of the tumour suppressor genes *Nf1* (N) and *Pten* (P) and concomitant expression of the constitutively active, oncogenic EGFR (E) variant *EGFRVIII* (hereafter termed NPE cells; **Figure 1A**). These NPE cells form aggressive GBM-like tumours upon transplantation and therefore model GBM-initiating stem cells. To investigate the function of ILK in NPE cells, we performed CRISPR/Cas9-mediated deletion of the *Ilk* gene in NPE cells (NPE^ILK−/−^) and re-expressed the ILK protein at endogenous levels (NPE^ILK-Rx^) in the same clonal population, thus creating paired isogenic NPE cells whose phenotypic differences can be attributed solely to expression of ILK (**Figure 1A**). ILK-deficient NPE^ILK−/−^ cells demonstrated loss of ILK’s core binding partners, PINCH-1 and ɑ-Parvin, as reported by others^25, 26^ (**Figure S1A**). Compared to ILK-expressing NPE and NPE^ILK-Rx^ cells, ILK-deficient NPE^ILK−/−^ cells were loosely attached to their substrate and more rounded in morphology, visibly displaying disorganised and poorly-polymerised actin and α-tubulin filaments; moreover, NPE^ILK−/−^ cells appeared to be more closely associated with each other in colonies (**Figure 1B**). These cytoskeletal alterations and morphologies were consistent in 5 NPE^ILK−/−^ clones and 2 guide RNAs (**Figure S1B, S1C**). Additionally, NPE^ILK−/−^ cells displayed reduced motility when compared to NPE and NPE^ILK-Rx^ cells in random single cell migration tracking assays (**Figure S1D; Supplementary Videos 1, 2 and 3**).

**Figure 1:**
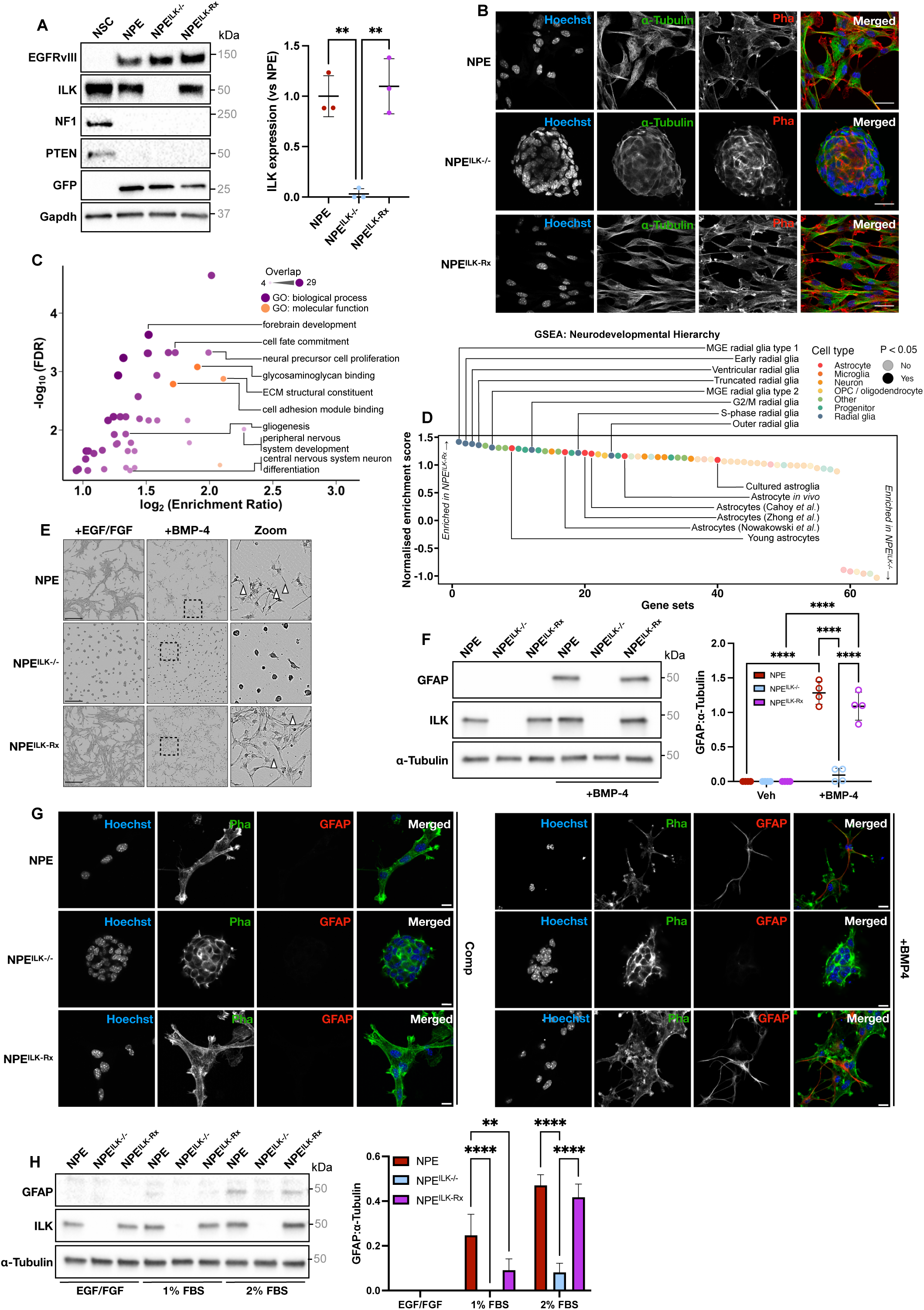
ILK controls GBM stem cell morphology and astrocyte differentiation. **(A)** Left: Western blot showing CRISPR/Cas9-mediated deletion of PTEN and NF1 and expression of GFP and EGFRVIII (“NPE”), ILK deletion (“NPE^ILK−/−^”), and ILK re-expression (“NPE^ILK-Rx^”) in NSCs. Right: Quantification of ILK expression in NPE, NPE^ILK−/−^ and NPE^ILK-Rx^ cells. (n = 3; ** *P* < 0.01, one-way ANOVA with Tukey’s multiple comparison test). **(B)** Immunofluorescence microscopy showing staining of nuclei (blue; Hoechst-33342), α-tubulin (green; α-tubulin antibody), and actin (red; Pha, phalloidin) in NPE, NPE^ILK−/−^ and NPE^ILK-Rx^ cells (n = 3 independent experiments). Scale bars = 20 µm. **(C)** Gene ontology (GO) term overrepresentation analysis of genes statistically significantly upregulated (FDR < 0.05; fold change > 2) in NPE^ILK-Rx^ *vs* NPE^ILK−/−^ cells. Shading intensity and point size indicate how many genes overlapped with reference terms; purple terms are related to biological processes and orange terms are related to molecular functions. Selected terms labelled. **(D)** Gene set enrichment analysis (GSEA) for 64 gene sets covering the neurodevelopmental hierarchy curated from published RNA-seq and scRNA-seq datasets. Gene sets are presented in descending order according to their normalised enrichment score, coloured according to gene set type, and shaded according to statistical significance (“MGE” = medial ganglionic eminence; “OPC” = oligodendrocyte precursor cell). **(E)** Phase-contrast micrographs of NPE, NPE^ILK−/−^ and NPE^ILK-Rx^ cells +/- bone morphogenetic protein-4 (BMP-4). White arrowheads mark processes observed in astrocyte-like, “stellate” morphology. Scale bars = 500 µm. **(F)** Left: Western blot showing expression of GFAP, an astrocyte marker, in NPE, NPE^ILK−/−^ and NPE^ILK-Rx^ +/- BMP-4 treatment for 24 h (n = 4). Right: Quantification of GFAP expression in NPE^ILK−/−^ and NPE^ILK-Rx^ +/- BMP-4 treatment for 24 h (n = 4; **** *P* < 0.001, two-way ANOVA with Šidák’s multiple comparison test). **(G)** Immunofluorescence microscopy staining nuclei (blue; Hoechst-33342), actin (green; Pha, phalloidin) and GFAP (red; GFAP antibody) in NPE, NPE^ILK−/−^ and NPE^ILK-Rx^ cells either without treatment (left), or following treatment with BMP-4 (right) (n = 3 independent experiments) (Scale bars = 20 µm). **(H)** Right: Representative Western blot showing GFAP expression in the indicated cell lines treated with increasing concentrations of foetal bovine serum (FBS). Left: Quantification of right (n = 3; \*\**P* < 0.01, \*\*\*\**P* < 0.005; two-way ANOVA with Tukey’s multiple comparison test).

As cell shape and adhesion are broadly correlated with transcriptional activity and stem cell biology^27, 28^, we asked whether ILK expression status affected transcription by performing 3’ mRNA sequencing (RNA-seq), hypothesising that ILK may regulate transcriptional programs associated with stem cell multipotency and differentiation. We detected 2,033 differentially-expressed genes between ILK deficient NPE^ILK−/−^ and ILK-expressing NPE^ILK-Rx^ cells, with expression of 1,025 genes increased in NPE^ILK−/−^ cells relative to NPE^ILK-Rx^ cells and 1,008 genes increased in NPE^ILK-Rx^ cells relative to NPE^ILK−/−^ cells (fold change > 1.5 and FDR < 0.05; **Figure S2A**). We performed gene ontology (GO) term analysis on 421 genes that were robustly upregulated in ILK-expressing NPE^ILK-Rx^ cells relative to NPE^ILK−/−^ cells (fold change > 2 and FDR < 0.05), and observed significant overrepresentation of biological process terms pertaining to neural lineage specification and nervous system development, including *cell fate commitment*, *gliogenesis* and *neural precursor cell proliferation* (**Figure 1C**). Consistent with ILK’s known functions as a mediator of integrin-ECM at focal adhesions and downstream signalling^7, 29^, we also noted over-representation of molecular function terms related to cell-ECM interaction including *cell adhesion module binding* and *extracellular matrix structural constituent* (**Figure 1C**). We did not observe overrepresentation of any biological process terms relating to NSCs or neurodevelopment when performing the same analysis on 479 robustly upregulated genes in ILK-deficient NPE^ILK−/−^ cells (**Figure S2B**). These data imply that ILK has both expected and previously unknown transcriptional effects in NPE cells, including effects pertaining to stem cell plasticity and fate. Additionally, noting that ILK depletion correlated with morphological change in NPE^ILK−/−^ cells resulting in a rounded and less mesenchymal-like morphology, we performed gene set enrichment analysis (GSEA) on a 194-member consensus gene set of genes regulating epithelial-to-mesenchymal transition (EMT)^30^. We found that the EMT gene set was significantly enriched in ILK-expressing NPE^ILK-Rx^ (**Figure S2C**), with 38 members (19.6%) of the gene set significantly upregulated (**Figure S2D**). Western blotting for the mesenchymal markers N-Cadherin, CD44 and MMP14^31, 32^ revealed significant depletion of each of these in NPE^ILK−/−^ cells *vs* parental NPE cells and NPE^ILK-Rx^ cells (**Figure S2E**). Finally, since mesenchymal phenotype is associated with invasion in many cancer types including GBM^33^, we performed 3-dimensional invasion assays by embedding NPE^ILK−/−^ or NPE^ILK-Rx^ neurospheres into Matrigel, a solubilised basement membrane preparation, and quantified their invasion over time. We observed inhibited invasion over time in NPE^ILK−/−^ cells relative to NPE^ILK-Rx^ cells (**Figure S2F, Supplementary Video 4, Supplementary Video 5**), correlating with loss of actin filament organisation and reduction of mesenchymal-like gene expression. These data indicate that ILK promotes the mesenchymal state and invasion in our GBM stem cell model.

Since the over-represented GO terms in NPE^ILK-Rx^ cells indicated a role for ILK in NSC plasticity and neurodevelopment (**Figure 1C**), we constructed a curated panel of 64 literature-reported gene sets sourced from bulk and single-cell RNA-Seq (sc-RNAseq) assays^34–37^. These describe diverse cell lineages in the neurodevelopmental hierarchy including radial glia, astrocytes, neurons, oligodendrocytes, progenitor cells, microglia, and diverse additional cell types (“other”) which are either uncharacterised^37^, or which are present in the brain but are not of neural lineage (such as endothelial cells and pericytes) (**Supplementary Table 1**). Differential GSEA between NPE^ILK−/−^ and NPE^ILK-Rx^ cells revealed that ILK-expressing NPE cells are enriched for a significant proportion (34 of 64; 53%) of the gene sets, which were most prominently astrocyte (6 of 9; 67%) and oligodendrocyte precursor cell (OPC; 2 of 2) linked but not mature oligodendrocyte (0 of 4) linked. Notably, each of the 8 tested radial glia gene sets were enriched in ILK-expressing NPE^ILK-Rx^ cells vs NPE^ILK−/−^ cells (**Figure 1D**), and included consensus radial glia markers such as *Blbp* (*Fabp7*), *Nestin, Tenascin-C* and *Hes1*^38, 39^ (**Figure S3A**): this finding is important given that the NPE cells most closely resemble radial glia^15, 23^, an NSC subtype which emerges mid-gestation and gives rise to diverse neuronal and glial lineages including astrocytes^40–42^. These data reveal a novel role for ILK in transcriptionally defining transformed radial glia identity.

### ILK controls GBM stem cell plasticity and transcriptional response to an astrocyte differentiation cue

GBM stem cells transition between different developmental states through differentiation processes, cell cycle changes, or responses to the tumour microenvironment such as altered cytokines or hypoxia^6, 15, 43^. In patient tumours these processes generate substantial transcriptional and phenotypic heterogeneity^44^. Since ILK loss resulted in the depletion of transcriptional signatures that describe radial glia (**Figure 1D**), plastic cells that give rise to diverse lineages represented both in the healthy brain and in GBM, we asked whether ILK-deficient NPE^ILK−/−^ cells could respond to BMP-4, which is cytostatic and is known to induce a non-proliferative astrocyte differentiation state^15, 22, 23^ that is positive for glial fibrillary acidic protein (GFAP), a marker of astrocyte-like cells^45, 46^. BMP-4-treated NPE and NPE^ILK-Rx^ (NPE-BMP and NPE^ILK-Rx^BMP cells) cells adopted a multi-process, “stellate” morphology characteristic of astrocytes, whereas ILK-deficient NPE^ILK−/−^ cells treated with BMP-4 (NPE^ILK−/−^BMP) retained their rounded, epithelial-like morphology (**Figure 1E**). Western blotting for GFAP revealed induction of expression in NPE-BMP and NPE^ILK-Rx^BMP cells, but not in NPE, NPE^ILK−/−^ and NPE^ILK-Rx^ (untreated control) cells or NPE^ILK−/−^ BMP cells (**Figure 1F**). Additionally, only NPE-BMP and NPE^ILK-Rx^BMP cells were GFAP-positive with stellate morphology in immunofluorescence assays (**Figure 1G**), indicating that loss of ILK prohibits transition from the NSC/radial glia-like phenotype to an GFAP-expressing astrocyte-like state in NPE cells.

Finally, to assess whether ILK-dependent GBM stem cell-to-astrocyte state transition strictly limited to BMP-4-induced effects, we treated NPE, NPE^ILK−/−^ and NPE^ILK-Rx^ cells with increasing doses of serum, known stimulator of NSC and GBM stem cell differentiation to astrocytes^22, 23^. We found that ILK was required for efficient GFAP expression at each serum concentration tested (**Figure 1H**), with ILK-expressing NPE and NPE^ILK-Rx^ cells expressing more GFAP than their ILK-deficient NPE^ILK−/−^ counterparts, indicating that ILK loss inhibits astrocyte differentiation in multiple contexts. These data suggest a function for ILK-mediated signalling in controlling stem cell-to-astrocyte transitions under multiple conditions in our GBM stem cells.

We next explored the transcriptional differences between NPE, NPE^ILK−/−^ and NPE^ILK-Rx^ cells following treatment with BMP-4 using RNA-seq. Pairwise hierarchical clustering analyses between each of the cell lines and treatment conditions revealed a distinct transcriptional cluster comprised exclusively of ILK-expressing NPE-BMP and NPE^ILK-Rx^BMP cells, whereas each of the other cells and conditions segregated into a single cluster (**Figure 2A**). These data indicate that ILK-deficient NPE^ILK−/−^BMP cells more closely resembled their untreated ILK-expressing counterparts than BMP-4-treated ILK-expressing NPE-BMP and NPE^ILK-Rx^BMP cells, suggesting that ILK loss resulted in a suppressed transcriptional response to BMP-4.

**Figure 2:**
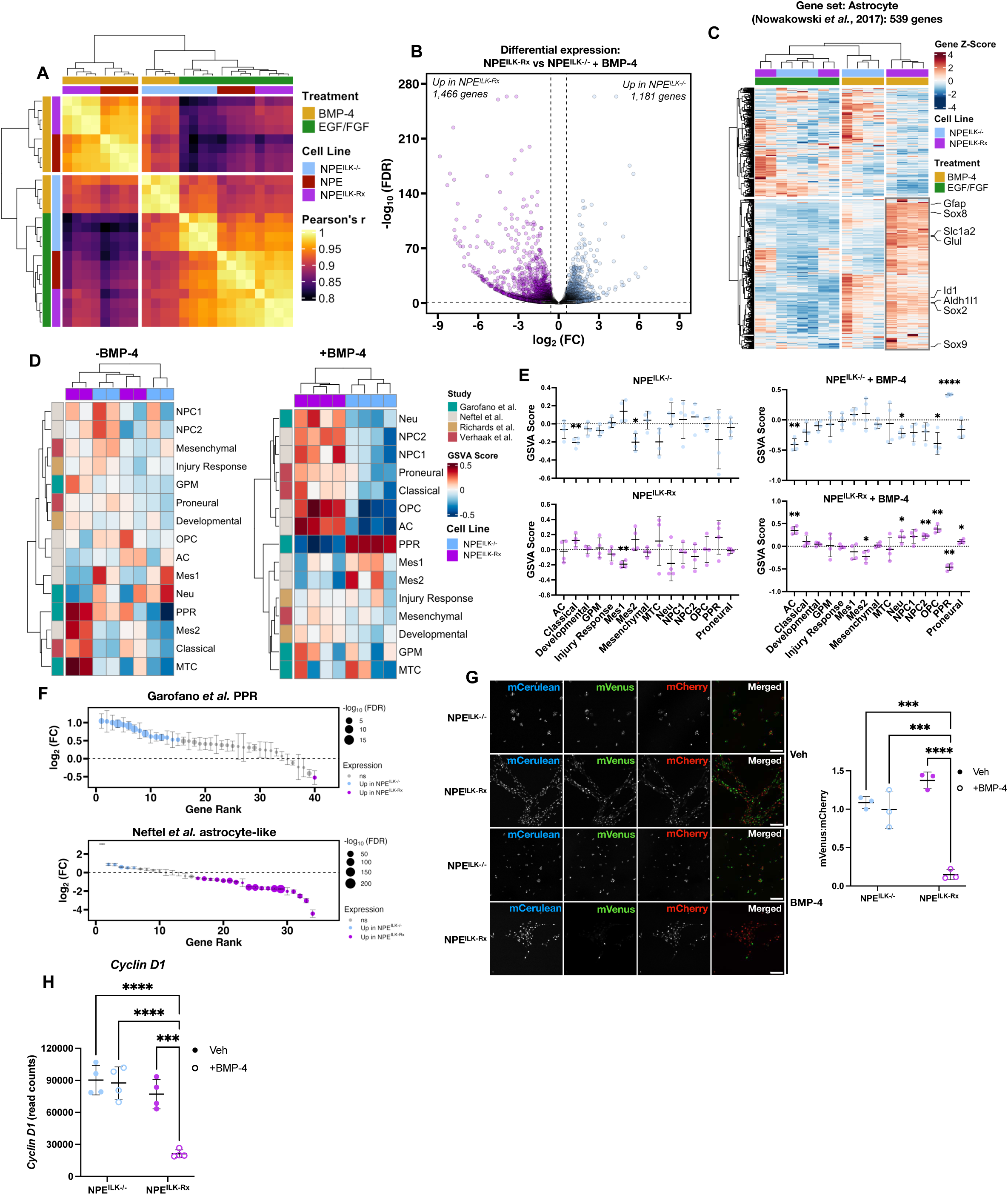
ILK controls transcription and GBM cell state transitions in response to BMP-4. **(A)** Heatmap showing correlation coefficients between NPE, NPE^ILK−/−^ and NPE^ILK-Rx^ transcriptomes +/- BMP-4 treatment for 24 h. Annotations indicate cell line and treatment; brighter colours indicate greater correlation between samples (*r*: Pearson correlation coefficient). **(B)** Volcano plot showing gene expression changes in NPE^ILK−/−^ vs NPE^ILK-Rx^ cells following treatment with BMP-4 (n = 4). Purple points are genes enriched in NPE^ILK-Rx^ cells; blue points are genes enriched in NPE^ILK−/−^ cells (Cutoffs of 1.5-fold expression change and false discovery rate (FDR)-adjusted *P* value < 0.05, Student’s *t*-test). **(C)** Heatmap showing differential expression of astrocyte genes in NPE^ILK−/−^ vs NPE^ILK-Rx^ cells either +/- BMP-4 treatment for 24 h. Annotations indicate cell line and treatment; red indicates relative enrichment, blue indicates relative depletion. Selected consensus astrocyte marker genes labelled. **(D)** Heatmaps showing enrichment of each of the investigated GBM cell states, as determined using gene set variation analysis (GSVA), in NPE^ILK−/−^ and NPE^ILK-Rx^ cells, either without treatment (left) or following treatment with BMP-4 (right); red indicates relative enrichment, blue indicates relative depletion. Annotations indicate cell line and the study that initially reported each GBM cell state. **(E)** Quantification of (D). Expression of each GBM gene set in NPE^ILK−/−^ and NPE^ILK-Rx^ cells (top and bottom), either without treatment (left) or following treatment with BMP-4 (right) (n = 4; * *P* < 0.05; ** *P* < 0.01; **** *P* < 0.001; One-sample *t*-test *vs* a hypothetical mean of 0). **(F)** Expression of proliferative/progenitor (PPR) and astrocyte-like gene signatures in NPE^ILK−/−^ *vs* NPE^ILK-Rx^ cells treated with BMP-4 for 24 h. Coloured points indicate statistically significant enrichment (FDR < 0.05); point sizes indicate negative log_10_-transformed FDR values (n = 4). **(G)** Left: representative fluorescence micrographs showing expression of H2B-mCerulean (nuclei; blue), mVenus-hGeminin (S/G2/M-phase; green), and mCherry-Cdt1 (G1-phase; red) in NPE^ILK−/−^ and NPE^ILK-Rx^ cells, +/- BMP-4 treatment for 24 h (scale bar = 100 µm). Right: quantification of mVenus-hGeminin:mCherry-Cdt1 ratios in NPE^ILK−/−^ and NPE^ILK-Rx^ cells +/- BMP-4 treatment for 24 h (n = 3; * *P* < 0.05; ** *P* < 0.01; *** *P* < 0.005; **** *P* < 0.001, one-way ANOVA with Tukey’s multiple comparison test). **(H)** RNA expression (expressed as normalised read counts) of the *Ccnd1* genes in NPE^ILK−/−^ and NPE^ILK-Rx^ +/- BMP-4 treatment for 24 h (n = 4; * *P* < 0.05; ** *P* < 0.01; **** *P* < 0.001; two-way ANOVA with Šidák’s multiple comparison test).

We next performed differential gene expression analysis in BMP-4 treated cells, detecting 1,466 genes upregulated in ILK-expressing NPE^ILK-Rx^BMP cells and 1,181 genes upregulated in ILK-deficient NPE^ILK−/−^BMP cells (**Figure 2B**). Although GFAP is widely used as a marker of parenchymal astrocytes, its expression is detectable in other cell types within the astroglial lineage including outer radial glia and quiescent NSCs^46, 47^. Therefore, to determine the specific transcriptional state of the ILK-deficient and ILK-expressing cells treated with BMP-4, we examined expression of a 539-member astrocyte gene signature previously defined by scRNA-seq^37^. Hierarchical clustering revealed a module of 57.9% (312 of 539 genes) of the gene signature that was upregulated prominently in NPE^ILK-Rx^BMP cells, indicating robust ILK-dependent BMP-4 mediated astrocytic differentiation. This module included markers of both radial glia subsets and astrocytes (*Gfap*, *Aldh1l1* and *Slc1a2*), transcriptional regulators involved in astrocyte differentiation (*Id1*, *Sox8* and *Sox9*), glutamine synthase (*Glul*, a marker of differentiated astrocytes), and *Sox2*, which is elevated in quiescent cancer stem cells and mature astrocytes^46, 48–54^, each of which were enriched in BMP-4 treated NPE^ILK-Rx^ cells *vs* NPE^ILK−/−^ cells (**Figure 2C**, **S3B>**). These data demonstrate a dependency on ILK for the transcriptional response to the BMP-4 astrocytic transition cue, which correlates with its role in the maintenance of NSC and radial glia-like identity permissive for differentiation.

### ILK permits GBM stem cell plasticity and is required for cell state transition following BMP-4 treatment

A number of RNA-seq and scRNA-seq studies have reported that GBM tumours are comprised of transformed cells that transcriptionally align with and transition between one or more cell states, reflective of a GBM stem cell hierarchy^5, 6, 43^. Having established that ILK loss alters radial glia-like identity and accordingly prevents BMP-4-induced transition of NPE cells to an astrocyte-like state, we next set out to test whether any of the literature-reported transcriptional signatures that describe GBM stem cells in various developmental states were different between ILK-expressing NPE^ILK-Rx^ and ILK-deficient NPE^ILK−/−^ cells with and without BMP-4. We therefore performed gene set variation analysis (GSVA), a method for assessing the enrichment of gene signatures by comparing expression of genes within a gene set to those outside of it across groups of samples^55^, with a collection of 15 gene sets corresponding to GBM-relevant transcriptional states^6, 43, 56, 57^ (**Figure S4A**). ILK-expressing NPE^ILK-Rx^ and ILK-deficient NPE^ILK−/−^ cells demonstrated no differential enrichment of this collection of GBM transcriptional states, with hierarchical clustering showing no ILK-dependent separation between the cell lines (**Figure 2D**), and neither NPE^ILK-Rx^ nor NPE^ILK−/−^ cells featured significant enrichment of any GBM transcriptional states (**Figure 2E**). This observation is consistent with the stem-like state of NPE cells in culture and suggests that, with respect to GBM cell states, NPE cells represent a predominantly stem-like population that lacks lineage specification and is capable of generating diverse differentiated progeny^56^. In contrast, BMP-4-treated ILK-expressing NPE^ILK-Rx^BMP and ILK-deficient NPE^ILK−/−^BMP cells clustered distinctly from each other (**Figure 2D**), with NPE^ILK-Rx^BMP cells displaying a diverse range of GBM transcriptional states. These included the expected astrocyte-like GBM transcriptional state, as well as the Neu, NPC2, OPC and PN GBM states (**Figure 2E**, **S4B>**). This finding is consistent with the ability of GBM stem cells to transition to diverse cell states^5, 6^. However, NPE^ILK−/−^BMP cells failed to establish any of these cell states, instead demonstrating exclusive enrichment for a cell state that is reported to mark poorly-differentiated and proliferative progenitor cells (proliferative/progenitor; PPR)^56, 58^ (**Figure 2E**). This observation reflects the lack of GBM stem cell-to-astrocyte differentiation we observed in ILK-deficient NPE^ILK−/−^ cells (**Figure 1E, 1F**) treated with BMP-4. Differential expression analysis revealed significant (FDR < 0.05) enrichment of 35.7% (15 of 42) of PPR genes in NPE^ILK−/−^ cells and 52.7% (19 of 36) of astrocyte genes in NPE^ILK-Rx^ cells treated with BMP-4 (**Figure 2F**). We conclude that ILK is required for BMP-4-mediated transition to an astrocyte-like state and indicate that ILK-deficient NPE^ILK−/−^ cells are impaired in their ability to transition between cell states, and become ‘trapped’ in a progenitor-like cell state. ILK is therefore required for differentiation responses and associated transcriptional plasticity seen in GBM cells.

To functionally validate activity of the PPR gene signature in BMP-4-treated NPE^ILK−/−^ cells, we generated a tricistronic H2B-Cerulean-Fucci2a construct that combined the Fucci2a cell cycle reporter^59^ with a Histone H2B-Cerulean fusion. This permits tracking of H2B-Cerulean labelled nuclei through all cell cycle phases. Due to their reciprocal degradation during cell cycle progression, the Fucci2a probes allow image-based segmentation and quantification of cells in G1 phase of the cell cycle by mCherry-hCdt1 accumulation (red), and cells in S/G2/M phase by mVenus-hGeminin accumulation (green)^59^. Expression of H2B-Cerulean-Fucci2a ILK-deficient NPE^ILK−/−^ and ILK-expressing NPE^ILK-Rx^ cells and BMP-4 treatment revealed increased accumulation of NPE^ILK-Rx^BMP cells in the G1 phase of the cell cycle, consistent with the cytostatic effects of BMP-4-mediated astrocyte state transition. Conversely, ILK-deficient NPE^ILK−/−^BMP cells demonstrated no significant accumulation in G1, instead exhibiting proliferation persistence following exposure to BMP-4 (**Figure 2G**). Consistently, transcriptional expression *cyclin D1* (*Ccnd1*), an essential regulator of G1-S-phase transition, was significantly reduced in NPE^ILK-Rx^ BMP but not NPE^ILK−/−^BMP cells (**Figure 2H**). These data indicate that ILK is required for BMP-4-induced cytostatic effects and transcriptional response resulting in expression of astrocyte lineage markers, and that ILK-deficient NPE^ILK−/−^ cells are sequestered in a proliferative/progenitor state that functionally describes their lack of state transition in response to BMP-4.

### An ILK-STAT3 pathway controls astrocyte-like state transition in GBM stem cells

To deduce the mechanism through which ILK permits BMP-4 response, we used reverse-phase protein array (RPPA), a functional proteomic assay for parallel analysis of many signalling proteins and/or their phosphorylation states, to compare signalling in ILK-expressing NPE-BMP cells with ILK-deficient NPE^ILK−/−^BMP cells. We found that treatment with BMP-4 induced significant changes to expression or phosphorylation of 12.5% (15 of 120) of the investigated proteins, with a further 11.6% (14 of 120) of changes induced by ILK loss independent of BMP-4 treatment (**Figure 3A**). These were broadly related to protein kinases and specific phosphorylation events, and included ILK-dependent induction of phospho- (p)Smad1/5, pSTAT3, pAkt and pEphA2 (**Figure 3A**). As BMP family members canonically signal via the Smad1/5 pathway^60^, we expected that ILK-deficient NPE^ILK−/−^BMP cells would display reduced Smad1/5 phosphorylation when compared to NPE-BMP cells. However, we observed the opposite, with a significant increase in Smad1/5 signalling in ILK-deficient NPE^ILK−/−^BMP cells when compared to ILK-proficient NPE-BMP cells that was not accompanied by an increase in Smad2/3 signalling, which has previously been linked to maintenance of the pluripotent state stem cell state^61, 62^ (**Figure 3A**, **S5A>**). BMP-4 can also signal via janus kinase (JAK) and signal transducer and activation of transcription (STAT)^63^, and dependence upon STAT3 in differentiation of non-transformed astrocytes has been reported^64, 65^. Consistently, we observed enhanced phosphorylation of STAT3 on residue Y705 (pSTAT3 Y705) in ILK-expressing NPE-BMP cells when compared to ILK-deficient NPE^ILK−/−^BMP cells (**Figure 3A**). We found no significant differences in phosphorylation of STAT3 S727 which, in addition to Y705, has been reported to regulate stem cell proliferation and pluripotency^66^ (**Figure S5B**). We confirmed that ILK was required induction of STAT3 Y705 phosphorylation in BMP-4 treated cells using western blotting, observing a concomitant robust increase in GFAP expression and STAT3 Y705 phosphorylation in BMP-4-treated ILK-proficient NPE-BMP and NPE^ILK-Rx^BMP cells but not in ILK-deficient NPE^ILK−/−^BMP cells, demonstrating that STAT3 Y705 phosphorylation was, like induction of GFAP, ILK-dependent (**Figure 3B**). This identified a new signalling pathway between ILK and STAT3 phosphorylation at residue Y705 that correlates with BMP-4-mediated pro-astrocytic effects.

**Figure 3:**
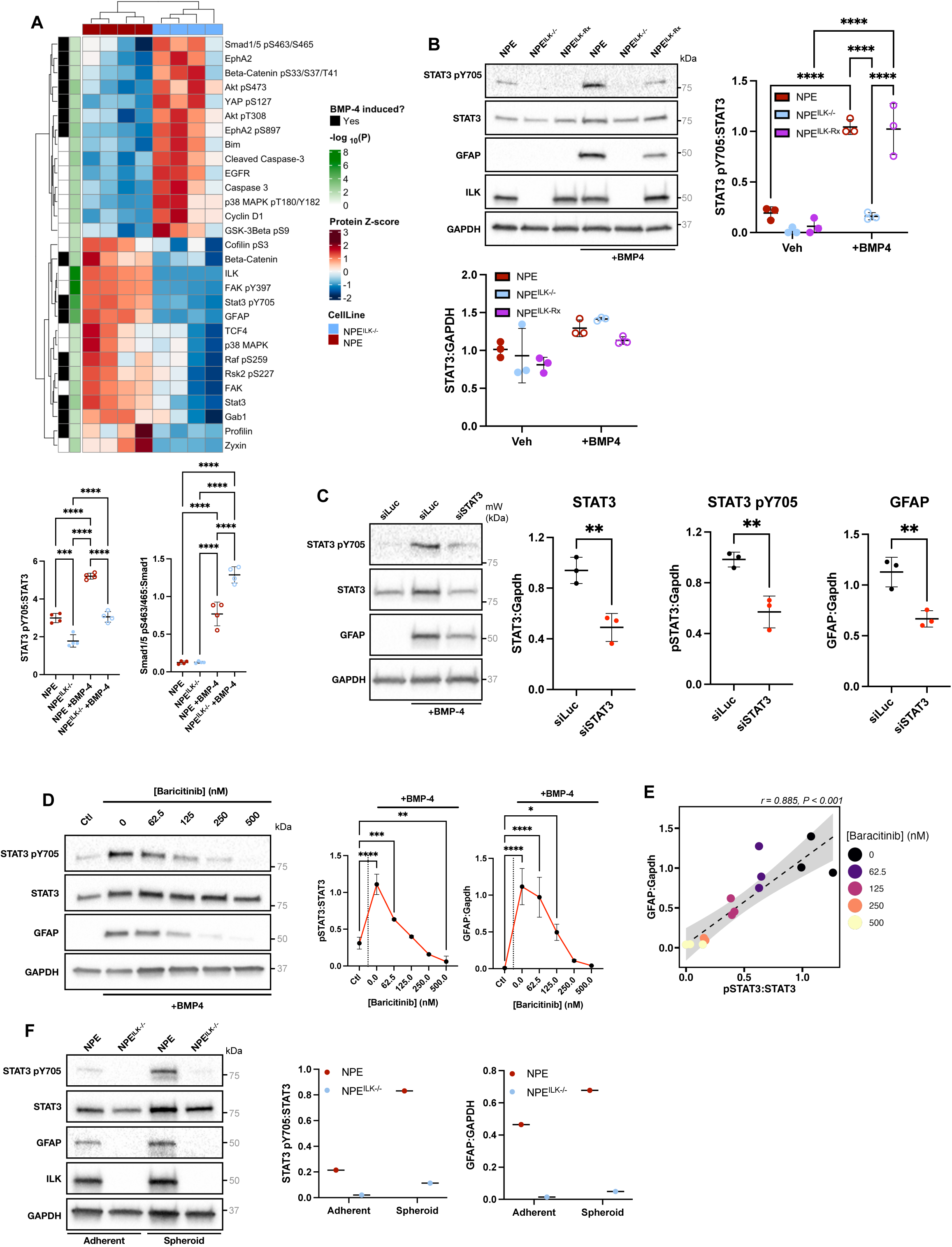
An ILK-STAT3 pathway controls GBM stem cell-to-astrocyte state transition. **(A)** Top: Heatmap showing statistically significant (*P* < 0.05; Student’s *t-*test) differential enrichment of the indicated proteins and phosphorylation sites in NPE *vs* NPE^ILK−/−^ cells as determined by reverse phase protein array (RPPA). Red indicates relative enrichment, blue indicates relative depletion; annotations indicate cell line, statistical significance level and changes that were induced in cells treated with BMP-4 but not untreated cells. Bottom: phosphorylation of the indicated factors normalised to total expression of each factor in NPE *vs* NPE^ILK−/−^ cells +/- BMP-4 treatment for 24 h (n = 4; *** *P* < 0.005; **** *P* < 0.001; one-way ANOVA with Tukey’s multiple comparison test). **(B)** Top left: Western blot showing phosphorylation and expression of STAT3 in NPE, NPE^ILK−/−^ and NPE^ILK-Rx^ cells +/- BMP-4 treatment for 24 h. Top right, bottom left: quantification of the indicated factors in NPE, NPE^ILK−/−^ and NPE^ILK-Rx^ cells +/- BMP-4 treatment for 24 h (n = 3; ** *P* < 0.01; **** *P* < 0.001; two-way ANOVA with Šidák’s multiple comparison test). **(C)** Western blot expression and phosphorylation of the indicated factors in NPE cells treated with small interfering (si) RNA targeting the *Luc* (negative control) gene in untreated cells, or siRNA targeting either *Luc or Stat3* in BMP4 treated cells. Right: Quantification of GFAP expression and STAT3 expression and phosphorylation in BMP4 treated NPE cells (n = 3; ** *P* < 0.01; Student’s *t*-test). **(D)** Left: Western blot showing GFAP expression and phosphorylation and expression of STAT3 following treatment of NPE cells with BMP-4 and the indicated doses of the JAK1/JAK2 inhibitor baricitinib (n = 3). Right: quantification of left, showing pSTAT3:STAT3 and GFAP:GAPDH ratios in NPE cells treated with BMP-4 and increasing doses of baricitinib (n = 3; * *P* < 0.05; ** *P* < 0.01; *** *P* < 0.005; **** *P* < 0.001, one-way ANOVA with Tukey’s multiple comparison test). **(E)** Model describing the relationship between STAT3 pY705:STAT3 and GFAP: GAPDH in NPE cells treated with BMP-4 and baricitinib. Black dashed trendline shown; shaded grey area indicates standard error; colours indicate doses of baricitinib (*r*: Pearson correlation coefficient). **(F)** Left: Western blot showing GFAP expression and phosphorylation and expression of STAT3 in NPE and NPE^ILK−/−^ cells and neurospheres. Right: Quantification of left (n = 1).

We next addressed whether STAT3 was required for the NSC-to-astrocytic state transition by performing short interfering(si)-RNA experiments in NPE cells, using an siRNA targeting either *Stat3* (siSTAT3) or *luciferase* (siLuc) as a control. Although we only observed partial (48.9%) *Stat3* knockdown with siSTAT3 in NPE-BMP cells, we found a concomitant reduction of both STAT3 pY705 (42.0%) and GFAP (40.9%) levels, consistent with a causal link between STAT3 signalling and GFAP expression (**Figure 3C**). We also treated ILK-proficient NPE-BMP cells with a concentration dose-response series of baricitinib, a clinically-approved inhibitor of the STAT3 kinases JAK1 and JAK2^67^. We observed inhibition of STAT3 Y705 phosphorylation, starting at a baricitinib dose of 62.5 nM and increasing to maximal inhibition at 250 nM, which correlated with reduction of GFAP expression in NPE-BMP cells (**Figure 3D**). Quantification of STAT3 pY705:STAT3 ratio and GFAP expression in BMP-4-treated NPE-BMP cells with increasing doses of baricitinib revealed a linear relationship (Pearson correlation coefficient = 0.885; **Figure 3E**), demonstrating strong association between STAT3 Y705 phosphorylation and GFAP expression. Taken together, these data imply that ILK-dependent STAT3 phosphorylation is causally linked to astrocytic differentiation. Finally, as ILK has adhesion-dependent functions^29^ and is part of the consensus adhesome^14^, we addressed whether ILK-dependent STAT3 Y705 phosphorylation was regulated by adhesion conditions. We therefore cultured ILK-proficient NPE and ILK-deficient NPE^ILK−/−^ cells in microplates coated with a low-attachment hydrogel to induce adhesion-independent neurosphere formation, removing the influence of direct attachment to laminin matrix employed in 2D culture. Western blotting revealed that neurosphere cultures retained ILK-dependent differences in STAT3 Y705 phosphorylation and GFAP expression, with NPE neurospheres producing increased levels of both GFAP and STAT3 pY705 in 3D cultures even without treatment with BMP-4 (**Figure 3F**). We also observed that in ILK-deficient NPE^ILK−/−^ neurospheres began to disaggregate from 4 days after initial formation (**Figure S6A, S6B**), with apparent lesions in the spheroid periphery and efflux of dead cells (as indicated by propidium iodide staining), from these lesions (**Figure S6C; Supplementary Videos 6, 7 and 8**). These data reveal an essential role for ILK-dependent STAT3 signalling that is distinct from ILK’s cell-ECM adhesion functions in astrocyte differentiation.

### The ILK-STAT3 pathway is active *in vivo* and correlates with GBM astrocyte-like and progenitor cell states

As discussed, GBM cells establish a range of diverse cell states *in vivo* and in patient tumours^43, 56^, and we found that ILK depletion results in loss of STAT3 signalling and failure to display BMP-4-mediated plasticity. To test the *in vivo* relevance of the ILK/STAT3 pathway, we injected mice intracranially with either NPE^ILK−/−^ or NPE^ILK-Rx^ cells and performed immunohistochemistry (IHC) staining for GFAP and STAT3 pY705 on excised brain slices. We found that NPE^ILK-Rx^ tumours stained strongly for GFAP (**Figure 4A**) and that this correlated with STAT3 pY705 staining; by comparison, NPE^ILK−/−^ cells displayed weaker GFAP staining and fewer cells displayed STAT3 pY705 positivity (**Figure 4A**). We additionally noted that NPE^ILK−/−^ cells exhibited reduced exfiltration from the injection site, correlating with our finding that ILK loss reduces cell motility and invasion (**Figure S1D**, **S2F>**, **S7A>**). To quantify ILK-dependent GFAP and STAT3 phosphorylation *in vivo*, we performed single-cell segmentation of IHC images, assigned cells to tumour or non-tumour groups based on GFP staining (as all NPE cells express GFP; **Figure S7A**), and quantified GFAP and STAT3 pY705 staining intensity per cell or per nucleus. We observed a pronounced increase in both GFAP and STAT3 pY705 staining relative to non-tumour brain in ILK-expressing NPE and NPE^ILK-Rx^ cells, but only modest changes in ILK-deficient NPE^ILK−/−^ cells (**Figure 4B**). ILK is therefore required for efficient GFAP expression and STAT3 Y705 phosphorylation *in vivo*, and an ILK-STAT3 pathway mediates astrocyte differentiation *in vitro* and correlates with GFAP expression *in vivo*. We assume that BMP-4, or other differentiating factors, are present in the brain microenvironment of tumour-bearing mice as previously reported^42^ and that these cause induction of STAT3 pY705 and GFAP in the presence of ILK. This defines a new pathway involving signalling from ILK to STAT3 pY705 phosphorylation, presumably via JAKs as suggested by effects of the JAK inhibitor baricitinib (**Figure 3D**), leading to induction of GFAP *in vitro* and *in vivo*.

**Figure 4:**
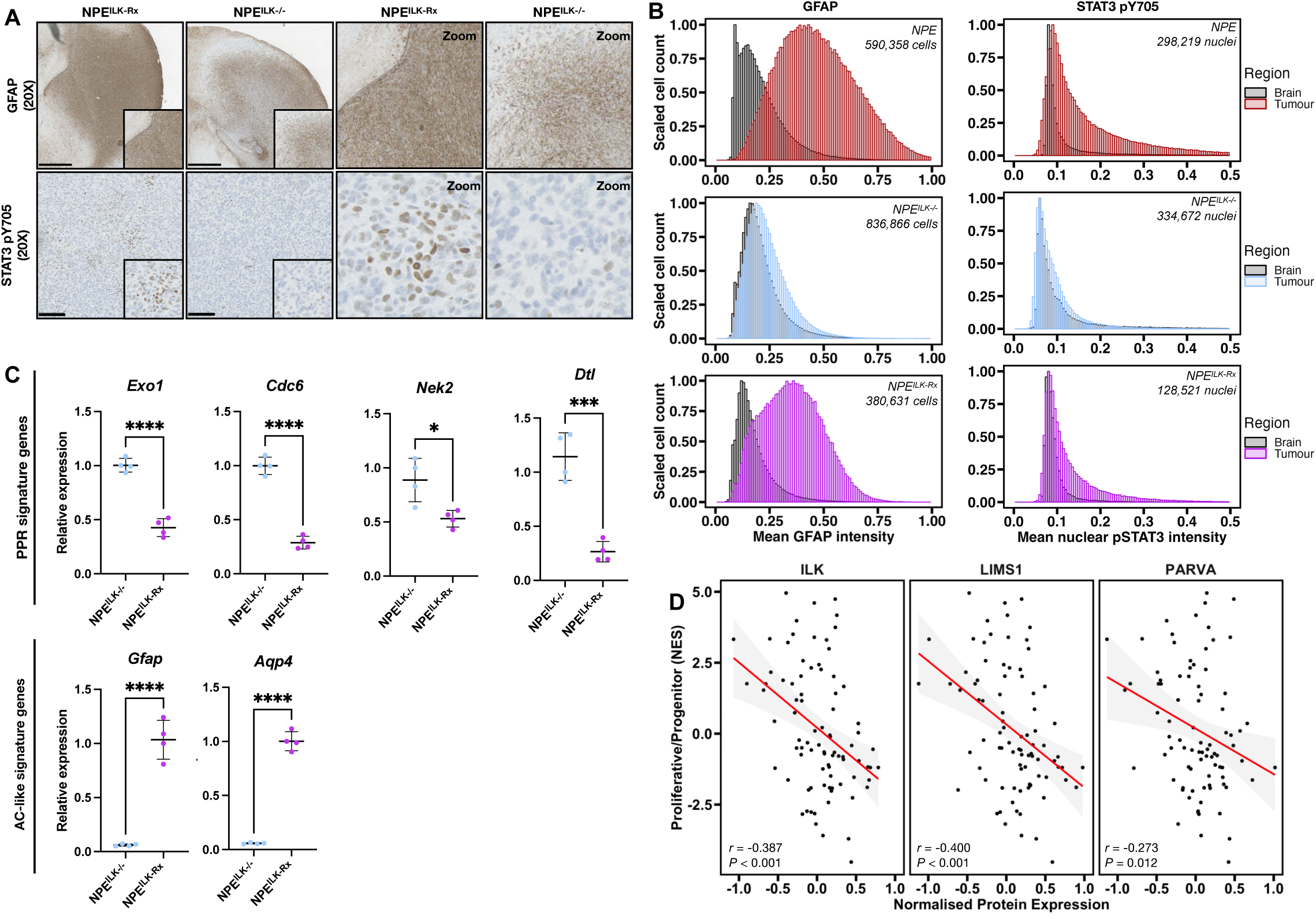
ILK-STAT3 pathway activity correlates with astrocyte and PPR cell state expression *in vivo* and in patients. **(A)** Representative immunohistochemistry images of GFAP and STAT3 pY705 staining in brain slices from mice intracranially injected with 5,000 NPE^ILK−/−^ or NPE^ILK-Rx^ cells (images representative of 4 mice). **(B)** Single cell quantification of either total GFAP staining or nuclear pSTAT3 Y705 staining in brain slices from mice intracranially injected with the indicated cell lines (quantifications representative of 4 mice). Histograms are coloured according to region as defined by GFP staining (tumour *vs* non-tumour brain; Figure S7). **(C)** Expression of the indicated PPR or astrocyte (AC)-like genes as assessed by qPCR in tumour slices from mice injected intracranially with NPE^ILK−/−^ or NPE^ILK-Rx^ cells (n = 4; * *P* < 0.05; *** *P* < 0.005; **** *P* < 0.001; Student’s *t*-test). **(D)** Correlation between protein expression of the indicated ILK / PINCH-1 (LIMS1) / α-Parvin (PARVA) complex proteins and PPR signature normalised enrichment scores (NES). Red trendline shown; shaded grey area indicates standard error (*r*: Pearson correlation coefficient).

Since we found that ILK-deficient NPE^ILK−/−^ cells predominantly express the PPR transcriptional signature *in vitro* and fail to respond to differentiation cues, we next asked whether ILK-deficient NPE^ILK−/−^ tumours were also enriched for the PPR transcriptional signature *in vivo*. Therefore, we extracted RNA from NPE^ILK−/−^ or NPE^ILK-Rx^ tumour slices and performed quantitative polymerase chain reaction (qPCR) to assay the expression of PPR genes that were robustly differentially-regulated in our RNA-seq dataset (**Figure 2D**; **Figure 2H**). We detected enhanced expression of the PPR-associated genes *Exo1*, *Cdc6*, *Nek2* and *Dtl* in NPE^ILK−/−^ when compared to ILK-expressing NPE^ILK-Rx^ tumours, in-keeping with ILK-dependent enrichment of the PPR signature *in vivo* and *in vitro* (**Figure 4C**, top panels). We also assessed expression of *Gfap* and *Aqp4*, the latter a predominantly astrocyte-associated gene and member of the astrocyte-like GBM gene signature, to validate whether NPE^ILK-Rx^ tumours contained astrocyte-like cells rather than GFAP-expressing radial glia^68^. We detected enhanced expression of *Aqp4* in ILK-expressing NPE^ILK-Rx^ tumours that was absent from ILK-deficient NPE^ILK−/−^ tumours (**Figure 4C**). Taken together, these data together imply that ILK is required to provide the signalling competence for transformed NSCs to transition into an astrocyte-like state *in vivo*, and that ILK loss results in the maintenance of a poorly-differentiated proliferative/progenitor signature, as we found *in vitro* (**Figure 2F**). This is likely a function of the new pathway we described whereby signalling from ILK to STAT3 pY705 regulates the transcription of specific differentiation-associated genes that leads, in turn, to plasticity in GBM tumours.

Finally, we asked whether our findings in our GBM model were reflected in patient tumours. We therefore made use of a recently-published multi-omic resource produced by the Clinical Proteomic Tumour Analysis Consortium^69^, which details proteomic and transcriptomic characterisation of 99 patient tumours, and integrated this data with recently-performed analysis of PPR transcriptional subtype enrichment in each of these patient samples^58^. We correlated protein expression of ILK, each of its binding partners, PINCH-1 (LIMS1) and α-Parvin (PARVA) in the IPP complex, and its integrin binding partners integrin β1 and integrin β3 (ITGB1 and ITGB3) with PPR signature scores in GBM patients; in each case, we observed a significant negative correlation (**Figure 4D**, **S7B>**). These data show that low ILK protein expression in patients correlates with PPR signature-rich tumours, consistent with our finding that ILK deficiency enriches for the PPR GBM cell state associated with suppressed STAT3 signalling and lack of plasticity, and suggest that our data in a mouse transformed NS cellular model is directly relevant to human GBM.

## Discussion

GBM displays substantial intratumoural heterogeneity that arises from GBM stem cells adopting diverse functional states and their inherent ability to switch between these^5, 6^. This plasticity that permits cell state transitions is controlled, at least in part, by the tumour microenvironment through mechanisms that are not well understood in GBM. Plasticity leads to challenges for anti-cancer therapeutics, with intratumoural heterogeneity and cell state transitions contributing to intrinsic and acquired drug resistance. Although detailed studies have described GBM stem cell plasticity and multiple transcriptional cell states have been defined^5, 6, 43, 56^, few critical regulators of GBM cell plasticity that may represent actionable therapeutic targets have been identified. This is important as their inhibition may ‘trap’ GBM stem cells in a single, or predominant, state, suppressing plasticity-mediated heterogeneity and leading to vulnerabilities that may be exploited for therapy.

Here we show that ILK, a protein that is best understood to function at integrin adhesion complexes as part of the ILK-Pinch-Parvin (IPP) complex^29^, has a novel and unexpected function in controlling GBM stem cell state and plasticity in a genetically-defined GBM stem cell model^15^. This role for ILK fits with the notion that extracellular matrix interactions, mediated by integrins and their downstream effectors, contribute to plasticity. ILK therefore has dual roles in GBM, controlling both adhesion and motility and associated malignant behaviours, and enabling signalling pathways that support differentiation and plasticity. We describe a novel ILK-STAT3 pathway that is required for transition of GBM stem cells to an astrocyte-like state in response to multiple stimuli (including anchorage-independent growth and both serum and BMP-4 treatment), and mediates the transition between cell states *in vitro* and *in vivo* when grown as tumours in the brains of recipient mice. Signalling via STAT3 has gained significant interest as a therapeutic target in cancers including GBM, and clinically-approved inhibitors of STAT3 upstream kinases are available. One of these is baricitinib, a JAK1/2 inhibitor that inhibits ILK-dependent phosphorylation of STAT3 Y705^67^. Our data therefore highlight a therapeutically-actionable mechanism of suppressing GBM stem cell plasticity by inhibiting ILK or ILK-dependent signalling to STAT3 that is likely via JAK1/2.

An outstanding question is how ILK deficiency contributes to loss of STAT3 Y705 phosphorylation. This is presumably indirect as ILK is a pseudokinase with no catalytic activity^7, 29, 70^; our data with baricitinib suggest that JAK1/2 is responsible for ILK-dependent phosphorylation of STAT3 that is linked to induction of differentiation and plasticity. One possibility is that loss of the adaptor functions of ILK may lead to mis-localisation of JAK1/2, normally localised to the BMP4-receptor in response to ligand. However, we confirmed that ILK-deficient cells retain their ability to activate signalling via the BMP receptor effectors Smad1/5 following BMP-4 exposure, indicating that receptor activity *per se* is not impaired (**Figure 3A**).

We found that loss of ILK gives rise to a stem cell population that is sequestered in a previously-defined proliferative/progenitor (PPR) transcriptional state^56, 58^ *in vivo* and *in vitro* when treated with BMP-4, a factor that normally induces cell cycle exit and astrocyte differentiation^23^. Interestingly, ILK has previously been implicated in suppressing NSC proliferation through its role in scaffolding PINCH-1 and Ras suppressor unit 1 (Rsu1), although we found that ILK depletion resulted in a retained proliferative phenotype as opposed to increased proliferation in our GBM stem cell model^71^. An important question is whether maintaining cells in a potentially proliferative state as a result of ILK loss may adversely lead to enhanced tumour growth. However, recent work has shown that PPR-like cells demonstrate a therapeutic vulnerability to a combination of radiotherapy and DNA-dependent protein kinase (DNA-PK) inhibitors, the latter as a result of their dependence on DNA-PK to repair DNA damage^58^. PPR-like cells are also poorly-invasive in comparison to other GBM cell states^58^, including astrocyte-like cells, which are enriched and undergo invasion at the tumour margins^72^. This is consistent with the lack of migration and invasion of ILK-deficient cells that we observed *in vitro* and *in vivo*, the latter from the injection site to outside the brain that only occurs when ILK is expressed (presumably via the needle track generated at the tumour cell injection site). Consistently, extracortical tumour growth is a reported feature of invasive GBM cells provided with a conduit for exfiltration from the brain^73^.

Given the phenotypic consequences of ILK loss in maintaining cells in a proliferative/progenitor transcriptional and functional state, we have contemplated whether inhibiting functioning of the ILK-STAT3 signalling pathway may best be exploited in the context of combination therapies. In favour of this, it is likely that abolition of the ability of GBM cells to transition exit the cell cycle (by maintaining a proliferative phenotype featuring cell cycle progression and retaining expression of *Cyclin D1*), which we show is lost upon ILK depletion in the context of BMP-4 treatment, may sensitise to treatment with agents such as temozolomide that preferentially targets cells in the G2/ M phase of the cell cycle^74^. We also found by RPPA-based functional proteomics that a number of signal transduction pathway components and phosphorylation events were increased in ILK-deficient cells, namely p38 MAP kinase pT180/Y182, GSK3b pS9, beta-catenin pS33/S37/T41, Akt pT308 and pS473, and others (**Figure 3A**). These likely represent compensatory survival mechanisms to cope with loss of ILK, suggesting that low doses of p38 inhibitors, Wnt inhibitors or PI3-kinase-Akt pathway inhibitors may synergise with ILK-deficiency or loss of signalling downstream. We also note enhanced expression of Caspase-3 and cleaved Caspase-3 upon ILK depletion (**Figure 3A**), implying that the cells may be primed for apoptosis. These possibilities, together with the proposed sensitivity of PPR-state to DNA-PK inhibition^58^, suggest that ILK-dependent vulnerabilities featuring prevention of cell cycle exit and reduction in plasticity will provide opportunities for combination therapies. Future work will strive to identify potential therapeutic combinations via co-inhibition with blockade of ILK-mediated GBM stem cell plasticity. To this end, we are developing small molecule agents that bind the ILK pseudo-active site, which we predict will impair formation of the IPP complex and signalling downstream to include in such combination inhibitor studies.

In addition to blockade of differentiation and the consequent ‘locking’ of transformed NSCs predominantly in a PPR-like transcriptional state, we found that ILK deficiency also profoundly affects cellular morphology and cytoskeletal organisation. The capacity of ILK-deficient cells to migrate and invade is significantly impaired as a result. Moreover, ILK is required for maintenance of a mesenchymal-like phenotype that is associated with efficient migration and invasion *in vitro*; this is perhaps unsurprising, given the known role for ILK at focal adhesion-based integrin-ECM contacts. Interestingly, STAT3 (in association with C/EBPβ) has been reported as a master regulator of mesenchymal phenotype in GBM^33^ and gliogenesis in the central nervous system during development^64, 65^. It is therefore likely that the apparently distinct morphological and differentiation/plasticity roles for ILK in transformed NSC model of GBM are co-regulated by the novel signalling ILK-STAT3 pathway we describe here, highlighting a key role for ILK in integrating malignancy-associated STAT3 signalling.

In summary, ILK has an unexpected role in rendering GBM cells permissive for cell cycle exit, differentiation and plasticity leading to diverse transcriptional cell states in tumour cell populations, while simultaneously promoting maintenance of mesenchymal-like morphology that is associated with migratory and invasive potential. This is in keeping with our finding that in patients, low expression of the ILK protein and its binding partners in the IPP complex correlates with the poorly-differentiated, poorly-invasive and constitutively cycling proliferative/progenitor cell state. We propose that the ILK-STAT3 pathway would be most effectively targeted with combination therapies via co-inhibition of ILK-STAT3 signalling and compensatory survival adaptations, and that co-inhibition of both cell plasticity and invasive potential may have beneficial effects in simultaneously stopping GBM from evolving away from therapeutic responses and blocking GBM cell invasion.

## Materials & Methods

### Cell culture and transfection

Murine NPE cells were generated using CRISPR/Cas9 mutagenesis from NSCs derived from the subventricular zone of adult C57BL/6-SCRM mice as previously described^15^. NPE cells were cultured in a humidified 37°C incubator at 5% CO_2_ in Dulbecco’s Modified Eagle Medium and Ham’s Nutrient Mixture F12 (DMEM/F12; Sigma-Aldrich, #D8437) supplemented with 1.45 g/L D-Glucose (Sigma-Aldrich, #G8644), 120 µg/mL bovine serum albumin (BSA) fraction V solution (Gibco, #15260-037), 100 µM β-mercaptoethanol (Gibco; #31350-010), 1X MEM non-essential amino acid (MEM-NEAA) solution (Gibco; #11140-035), 0.5X B-27 (Gibco; #17504-044) supplement, 0.5X N-2 supplement (Gibco; #15140-122), 10 ng/mL murine EGF (Peprotech; #315-09), 10 ng/mL human b-FGF (Peprotech; #100-18b), and 1 µg/mL laminin-I (R&D Systems; #3446-005-01). Cells were passaged every second day by dissociation with accutase (Sigma-Aldrich; #A6964) for 3 min at room temperature, centrifugation at 200 x *g* for 3 min, and resuspension in medium. Cells were routinely tested for mycoplasma infection.

For neurosphere assays, 2,000 cells were seeded into wells of 96-well U-bottom ultra-low attachment plates (Corning; #7007) and centrifuged at 200 x *g* for 5 min at room temperature. For invasion assays, neurospheres were cultured for 48 h and embedded into growth factor-reduced Matrigel® (Corning; #324320) diluted 1:1 in medium. Neurospheres were imaged every 3 hours for a period of 7 days on an IncuCyte® S3 live cell system (Satorius) in a humidified 37°C incubator at 5% CO_2_ using a 4X objective. Invasion was quantified using the Incucyte® Software (v2020C).

For single-cell tracking assays, 500 cells were plated into each well of an IncuCyte® ImageLock 96-well plate (Sartorius; #BA-04856) and imaged every 30 min for a period of 48 h using a 10X objective. Cell tracking measurements were made during the final 24 h using the ImageJ^75^ (v2.9.0/1.53t) Manual Tracking plugin.

For differentiation of NPE cells into astrocytes, cells were cultured in the absence of EGF and FGF and with the addition of either 10 ng/mL BMP-4 (Peprotech; #315-27) or 1% or 2% foetal bovine serum (Gibco; #10437-028) for 24 hours. Astrocytic differentiation was confirmed visually by adoption of typical “stellate” morphology and biochemically by evaluation of GFAP expression.

NPE cells were transfected by electroporation using a Lonza® Nucleofector 2b device and the Lonza® Mouse Neural Stem Cell Nucleofector Kit (Lonza; #VPG-1004) according to the manufacturer’s instructions using the T-030 pulse code.

### Plasmids

Plasmids for CRISPR/Cas9-mediated *Ilk* knockout were generated by cloning oligonucleotides encoding 2 sgRNAs targeting the *Ilk* gene (**1**) between the BbsI sites of the eSpCas9(1.1) plasmid^76^ (a gift from Feng Zhang; Addgene; #71814). For *Ilk* re-expression, total RNA was extracted from NPE cells and reverse-transcribed to produce a cDNA library according to the manufacturer’s protocols (Qiagen; #74104 and Thermo Scientific; #K1621). The *Ilk* gene was amplified from the cDNA library using primers introducing NotI and BamHI restriction sites into the 5’- and 3’-regions of the gene. The resulting construct was cloned between the NotI and BamHI sites of the pQCXIN plasmid (a kind gift from Toby Hurd). The piggyBac transposon plasmid pCAG-PBase was a kind gift from Richard Meehan. To generate H2B-Cerulean, mCerulean (Rizzo et al., 2004) was amplified from Cerulean^77^ (a gift from Dave Piston; Addgene #15214) by PCR and the product cut with KpnI, blunted, and cut with AgeI. In parallel, pH2B-GFP^78^ (a gift from Geoff Wahl; Addgene #11680) was cut with NotI, blunted, and cut with AgeI allowing subcloning of Cerulean to replace GFP. H2B-Cerulean-Fucci2a was generated by PCR amplification of H2B-Cerulean to introduce a porcine teschovirus-1 2A (P2A) self-cleaving peptide sequence and flanked by MluI and BssHII restriction sites allowing subcloning into the MluI site of pCDNA5-Fucci2a^59^. Finally, pB-GIP (a gift from Richard Meehan) was modified with a polylinker incorporating MluI and NruI sites allowing cloning of H2B-Cerulean-Fucci2a as an MluI/Eco53kI fragment to generate pB-H2B-Cerulean-Fucci2a.

### CRISPR/Cas9-mediated gene editing and stable cell line generation

1 x 10^6^ NPE cells were electroporated as described with 5 µg of either eSpCas9(1.1) or eSpCas9(1.1) with either *Ilk* sgRNA (**Supplementary Table 1**). This process was twice at 48 h intervals for a total of 3 rounds of electroporation. Cells were suspended in 5% BSA (Merck; #12659) in PBS and single cells were sorted using a BD FACSJazz system into wells of a 96-well cell culture plate. Cells were incubated in a humidified 37°C incubator at 5% CO_2_ until visible colonies were formed. Colonies were passaged into two wells of a 6-well plate, one of which was lysed directly in Laemmi buffer (50 mM Tris-HCl pH 6.8, 10% glycerol, 5% SDS, 5% β-mercaptoethanol, bromophenol blue), and processed for Western blotting to screen for ILK depletion. Where ILK depletion was detected, the remaining cells were passaged for further use.

### Antibodies

Antibodies used for Western blotting were anti-ILK (BD Biosciences; #611803, 1:2000), anti-STAT3 (Cell Signalling Technology; #12640, 1:1000), anti-STAT3 pY705 (Cell Signalling Technology; #9145, 1:1000), anti-GFAP (Cell Signalling Technology; #3670, 1:2000), anti-PINCH-1 (Cell Signalling Technology; #11890, 1:500), anti-ɑ-Parvin (Cell Signalling Technology; #8190, 1:1000), anti-NF1 (Bethyl; #A300-140A, 1:2000), anti-PTEN (Cell Signalling Technology; #9559, 1:1000), anti-GFP (Roche; #11814460001, 1:1000), anti-EGFR (Millipore; #04-290, 1:1000), anti-GAPDH (CST; #5174, 1:2000) and anti-ɑ-Tubulin (Cell Signalling Technology; #3873, 1:2000). Primary antibodies used for immunofluorescence microscopy were anti-ɑ-Tubulin (Cell Signalling Technology; #3873, 1:1000) and anti-GFAP (Cell Signalling Technology; #3670, 1:1000). Secondary antibodies used for immunofluorescence were Alexa Fluor 488-conjugated anti-rabbit IgG (Invitrogen; #A11008), Alexa Fluor 488-conjugated anti-mouse IgG (Invitrogen; #A21200), Alexa Fluor 568-conjugated anti-rabbit IgG (Invitrogen; #A11011) and Alexa Fluor 568-conjugated anti-mouse IgG (Invitrogen; #A11004, all 1:400). Antibodies used for immunohistochemistry were anti-GFAP (Cell Signalling Technology; #3670, 1:1000), anti-STAT3 pY705 (Cell Signalling Technology; #3760, 1:400) and anti-GFP (abcam; #ab290, 1:400). Antibodies used for reverse phase protein array (RPPA) are provided in **Supplementary Table 1**.

For re-expression of the *Ilk* gene in NPE^ILK−/−^ cells, 1 x 10^6^ cells were electroporated as described with 10 µg of ILK-pQCXIN. 48 h later, *Ilk*-expressing cells were selected by addition of 0.5 mg/mL G418 (Merck; #A1720). G418 selection was continued for 2 weeks to select only cells that had stably integrated the ILK-pQCXIN construct. Expression of ILK in G418-selected cells was confirmed by Western blotting. Selection was repeated every 3 months to ensure that the population retained ILK expression. For generation of H2B-Cerulean-Fucci2a-expressing cells, 1 x 10^6^ NPE, NPE^ILK−/−^ or NPE^ILK-Rx^ cells were electroporated as described with 5 µg pCAG-PBase and 5 µg pB-H2B-Cerulean-Fucci2a. 48 h later, H2B-Cerulean-Fucci2a-expressing cells were selected by addition of 1 µg/mL puromycin dihydrochloride (Thermo Scientific; #A1113803). Puromycin selection was continued for 2 weeks to select only cells that had stably integrated the H2B-Cerulean-Fucci2a construct. Expression of H2B-Cerulean-Fucci2a in puromycin-selected cells was confirmed by fluorescence microscopy using a Leica Stellaris 8 confocal microscope with the LasX software.

### Imaging and immunofluorescence microscopy

Coverslips were washed for 16 hours in 1 M hydrochloric acid at 65 °C before being rinsed twice with deionised water and stored in 70% ethanol. Prior to plating cells, coverslips were washed in PBS and coated with 1 µg / mL laminin-I (R&D Systems; #3446-005-01) at 37 °C for 1 hour. 5 x 10^4^ cells were plated on coated coverslips and allowed to adhere for 48 h. For imaging of fluorescent protein constructs, cells were fixed by direct addition of methanol-free formaldehyde (Thermo Scientific; #28908) to a final concentration of 4% and incubation at 37 °C for 15 min, washed twice with PBS, and mounted on coverslips using ProLong Glass Antifade Mountant (Invitrogen; #P36980). For immunofluorescence microscopy, cells were fixed by direct addition of formaldehyde (Sigma-Aldrich; #F1635) to a final concentration of 4% and incubation at 37 °C for 15 min. Cells were washed once with Tris-buffered saline (TBS) and incubated in 0.1 M glycine (Sigma-Aldrich; #G8898) in TBS for 10 minutes at room temperature to quench excess formaldehyde before an additional wash with TBS. Permeabilisation was performed with 0.1% Triton X-100 (Sigma-Aldrich; #T9284) in TBS for 5 minutes at room temperature. Cells were washed once in 0.05% Triton X-100 in TBS and blocked with 2% BSA in 0.1% Triton X-100 for 1 hour at room temperature. Primary antibodies were diluted in 2% BSA in 0.1% Triton X-100 in TBS and incubated with cells for 16 hours at 4 °C, followed by 3 washes with 0.1% Triton X-100 in TBS for 5 minutes at room temperature with gentle agitation. Secondary antibodies and, where used, Phalloidin-Atto647N (Sigma-Aldrich; #65906, working concentration 25 nM) were diluted in 2% BSA in 0.1% Triton X-100 and incubated with cells in the dark for 45 minutes at room temperature. Cells were washed a further 3 times in the dark in 0.05% Triton X-100 at room temperature for 5 minutes with gentle agitation, followed by a final rinse with deionised water. Cells were mounted with ProLong Glass Antifade Mountant with NucBlue (Invitrogen; #P36981) for nuclear staining. Cells were imaged on a Leica Stellaris 8 confocal microscope using a 405 nm diode (for imaging NucBlue and mCerulean) and a tuneable white light laser (tuned to 488 nm for imaging Alexa Fluor −488 and GFP, 515 nm for mVenus, 568 nm for Alexa Fluor −568, 594 nm for mCherry and 647 nm for Atto-647N) with either a 100X or 60X oil immersion objective, or a 20X air objective. Images were acquired using the Leica LasX software and processed using ImageJ^75^ (v2.9.0/1.53t).

### Western blotting

Cells were washed once with cold PBS and lysed in cold radioimmunoprecipitation assay buffer (RIPA buffer; 50 mM Tris-HCl pH 8, 150 mM NaCl, 1% Triton X-100, 0.5% sodium deoxycholate, 0.1% sodium dodecyl sulphate) supplemented with cOmplete™ ULTRA Protease Inhibitor (Roche; #5892953001) and PhosSTOP™ Phosphatase Inhibitor (Roche; #4906845001) cocktail tablets for 15 minutes at 4 °C with gentle agitation. Crude extracts were collected into microcentrifuge tubes and centrifuged at 4 °C for 15 minutes at 19,000 x*g*. Clarified protein-containing supernatants were collected and quantified using the Pierce™ BCA assay kit (Thermo Scientific; #23225) according to the manufacturer’s protocol. 15 µg protein was diluted in Laemmli buffer (final concentration 2% sodium dodechylsulphate, 10% glycerol, 50 mM Tris-HCl, pH6.8, 5% β-mercaptoethanol and 0.01% bromophenol blue) and heated to 95 °C for 5 minutes. Proteins were resolved by SDS-polyacrylamide gel electrophoresis using 4-15% Mini-PROTEAN® TGX™ gels (BioRad; #4561086) and transferred to nitrocellulose membranes (BioRad; #1704158) using the Trans-Blot Turbo semi-dry transfer system. Membranes were blocked by incubation with 5% BSA in 0.1% TWEEN® 20 (Millipore; #11332465001) in TBS (TBS-T) for 1 hour at room temperature with gentle agitation. Primary antibodies were diluted in 5% BSA in TBS-T and incubated with membranes for 16 hours at 4 °C with gentle agitation. Membranes were washed 3 times with TBS-T at room temperature for 15 minutes with agitation. Secondary antibodies were diluted in 5% BSA in TBS-T and incubated with membranes for 45 min at room temperature with agitation. Membranes were washed a further 3 times in TBS-T and bound antibodies were visualised using the Clarity Western ECL substrate (BioRad; #1705061) with a ChemiDoc MP system (BioRad). Images were collected and quantified using the BioRad ImageLab software (v6.1).

### Quantitative PCR

Cells were washed once with cold PBS and RNA was extracted using the RNeasy Mini Kit (Qiagen; #74104) with DNAse-I digestion according to the manufacturer’s instructions. Total RNA extracts were quantified using a Nanodrop™ 2000 spectrophotometer (Thermo Fisher; #ND-2000). mRNA was reverse-transcribed to cDNA by oligo-dT priming using the RevertAid Reverse Transcription Kit (Thermo Scientific; #K1691) according to the manufacturer’s instructions. For quantitative PCR, 25 ng cDNA was amplified using gene-specific primers (**Supplementary Table 1**) and the SYBR™ Select Master Mix kit (Applied Biosystems; #4472908) for a total of 35 cycles. Data were collected on a StepOnePlus™ PCR system (Applied Biosystems) using the StepOne™ Software (v2.3). Quantification was performed using the ΔΔCt method.

### Short interfering RNA and baricitinib treatment

To achieve short interfering RNA (siRNA)-mediated *Stat3* knockdown, we made use of a lipidoid nanoparticle-based strategy that suppresses transcript expression without the requirement for a transfection reagent. Briefly, *Stat3*-targeting siRNAs and control *Luc* siRNAs were packaged into three-component lipidod nanoparticles as previously described^79^. Sequences are reported in **Supplementary Table 1**. Cells were pre-treated with 1 µg/mL si*Luc* or si*Stat3*-loaded lipidoid nanoparticles for 24 h before addition of BMP-4, and for a further 24 h following addition of BMP-4.

For baricitinib treatment, cells were incubated with the indicated doses of baricitinib (APExBIO; #A4141) or DMSO. Cells were incubated with baricitinib for 24 h before addition of BMP-4, and for a further 24 h following addition of BMP-4.

### Animal studies

All procedures performed on mice were conducted in accordance with protocols approved by the Home Office in the United Kingdom under a project license for V.B. (PP7510272) at The University of Edinburgh. 10-week-old female CD-1/nude mice were injected intracranially with 10,000 cells suspended in 2 µL growth medium following anesthesia with isoflourane. For analgesia, buprenorphine was administered during surgery and carprofen was administered in water for a subsequent 48 h. Injections were performed 0.6 mm anterior and 1.5 mm lateral to the bregma at a depth of 2.5 mm. Tumour growth was monitored twice by weekly injection of 150 mg/kg luciferase and imaged using an IVIS® Lumina S5 system. Mice were sacrificed by cervical dislocation at 21 days post-injection. After sacrifice, brains were collected and immediately transferred to formalin for fixation for 16 hours.

### Immunohistochemistry

Brain samples from mice were cut into 4 coronal sections and embedded into paraffin blocks. Slices were cut at a thickness of 4 µm. Paraffin was removed from tissue sections by washing twice for 5 min in xylene and sections were rehydrated by washing for 3 minutes with 100%, 100%, 75% and 50% ethanol. Sections were rinsed in water before antigen retrieval by incubation in citrate buffer (0.825 M sodium citrate, 0.175 M citric acid; pH 6.0) at 100 °C for 7 minutes. Sections were rinsed with water and TBS-T twice for 5 minutes. Sections were treated with Dako REAL peroxidase block solution (Agilent; #S2023) for 5 min, rinsed in water, and treated for a further 10 min with serum-free protein block solution (Agilent; #X0909). Primary antibodies were diluted in antibody diluent (Agilent; #S3022) and incubated with sections for 16 hours at 4 °C. Samples were washed a further three times with TBS-T for 5 minutes and stained with DAKO EnVision-HRP rabbit/mouse (Agilent; #K5007) for 2 hours at room temperature. Samples were washed a further 3 times in TBS-T for 5 minutes and developed with DAB diluted at a 1:50 ratio in DAB-chromagen (Agilent; #K3468) until appearance of brown staining. Sections were rinsed with water and counterstained with Mayer’s hematoxylin solution (Agilent; #S3309) for 2.5 min at room temperature. Sections were rinsed twice with water and treated with Scott’s tap water (3.5 g/L sodium bicarbonate; 20 g/L magnesium sulphate) for 2 minutes before a final rinse with water. Sections were dehydrated by washing for 3 minutes with 50%, 75%, 100% and 100% ethanol and washed with xylene twice for 5 min at room temperature. Slides were mounted with DPX mountant (Sigma-Aldrich; #06522). Images of stained sections were acquired on a NanoZoomer slide scanner (Hamamatsu Photonics) and analysed with QuPath^80^ (v4.0.3) using default settings. Tumour and brain regions were segregated using GFP staining.

### Reverse phase protein array

Reverse phase protein array (RPPA) analysis was performed on nitrocellulose-coated slides as previously described^81^ by the Host and Tumour Profiling Unit at The University of Edinburgh. Briefly, cells were washed with PBS and lysed in RIPA buffer supplemented with cOmplete ULTRA protease inhibitor and PhosSTOP phosphatase inhibitor cocktails as described. Clarified lysates were serially diluted to produce a four-step doubling dilution series, which were spotted in technical triplicate onto nitrocellulose-coated slides (Grace Bio-Labs) under 70% humidity using an Aushon 2470 array platform (Aushon Biosystems). After hydration, slides were blocked using SuperBlock (TBS) blocking buffer (Thermo Fisher Scientific) and incubated with primary antibodies (all diluted 1:250 in SuperBlock; **Supplementary Table 1**). Bound antibodies were detected by incubation with anti-rabbit DyLight 800-conjugated secondary antibody (New England BioLabs). Slides were analysed using an InnoScan 710-IR scanner (Innopsys), and images were acquired at the highest gain without saturation of the fluorescence signal. The relative fluorescence intensity of each array feature was quantified using Mapix software (Innopsys). Normalised RPPA data are reported in **Supplementary Table 2**.

### RNA-sequencing

RNA-sequencing (RNA-seq) was performed by the Wellcome Trust Clinical Research Facility at The University of Edinburgh. RNA quality was assessed on an Agilent 2100 Electrophoresis Bioanalyser instrument (Agilent; #G2939AA) and an RNA 6000 NanoChip (Agilent; #5067-1511). RNA was quantified using a Qubit 2.0 Fluorometer (Thermo Scientific; #Q32866) and the Qubit RNA Broad Range assay kit (Thermo Scientific; #Q10210). Lack of DNA contamination was confirmed using the Qubit dsDNA HS assay kit (Thermo Scientific; #Q32854). Libraries were prepared from 110 ng of each RNA sample using the QuantSeq 3’ mRNA-Seq Library Prep Kit (FWD) for Illumina (Lexogen; #015) according to the manufacturer’s protocol. Library generation was initiated using oligo-dT priming with Illumina-compatible linker sequences. After first-strand synthesis, RNA was removed and second-strand synthesis was performed using random priming. Following synthesis, magnetic bead-based purification was performed before cDNA libraries were amplified for 17 cycles, introducing cluster generation and index sequences to allow multiplexing. Libraries were quantified using the Qubit dsDNA HS assay kit and assayed for quality and fragment size using a Bioanalyser with the DNA HS Kit (Agilent; #5067-4626). Sequencing was performed on a NextSeq 2000 platform (Illumina; #20038897) using NextSeq 1000/2000 P2 Reagents for 100 cycles (Illumina; #20046811). Libraries were combined in an equimolar pool and run on a P2 flow cell. Base calling was performed using the NextSeq 1000/2000 Control Software (v1.4.1.39716) and data were analysed using the QuantSeq data analysis pipeline integrated on the BlueBee genomics analysis platform. Read counts are reported in **Supplementary Table 2**.

### Computational methods

For RNA-seq analysis, read counts were normalised, low read counts (fewer than 10 reads across all samples) filtered, and differential expression analysis performed using DESeq2^82^ (v1.38.3). Multiple testing was accounted for using Benjamini-Hochberg correction to calculate false discovery rate (FDR).

For gene ontology (GO) term analysis, statistically significantly differentially-expressed genes (with cutoffs of FDR < 0.05 and fold change < 2) were submitted to the WebGestalt gene set analysis toolkit^83^ (https://www.webgestalt.org) using the Biological Process (noRedundant) functional database to reduce redundancy amongst overrepresented processes with a significance level cutoff of FDR < 0.05.

For gene set enrichment analysis (GSEA) and gene set variation analysis (GSVA), gene sets were assembled from reported signatures that describe GBM cell states and neurodevelopmental lineages^6, 34–37, 43, 56, 57^ and, where necessary, human gene names were converted to mouse gene names using the biomaRt R package (v2.54.1). GSEA was performed using the GSEA software^84^ (v4.3.2) on expression datasets containing normalised read counts filtered for only genes with counts of > 10 over all samples. The Signal2Noise metric was used for gene ranking. For GSVA, the GSVA R package^55^ (v1.46.0) was used on expression datasets (as filtered for GSEA) using default parameters.

Where R packages were used for analysis, this was performed using R (v4.2.2) and the RStudio Integrated Desktop Environment (2023.06.0+421). Data were plotted using GraphPad Prism 9 (v9.5.1), the ggplot2 package^85^ (v3.4.2), or the ComplexHeatmap package^86^ (v2.14.0).

## Supporting information

Supplementary Video 1: NPE migration

Supplementary Video 2: NPE ILK-/- Migration

Supplementary Video 3: NPE ILK-Rx migration

Supplementary Video 4: NPE ILK-/- invasion

Supplementary Video 5: NPE ILK-Rx invasion

Supplementary Video 6: NPE spheroid

Supplementary Video 7: NPE ILK-/- spheroid

Supplementary Video 8: NPE ILK-Rx spheroid

## Acknowledgements

We are grateful to core services staff at the Institute of Genetics and Cancer for their expertise and contributions to this work. Specifically, we thank Ann Wheeler and Matthew Pearson at the Advanced Imaging Resource for imaging support; Kenneth McLeod and Alison Munro at the Host and Tumour Profiling Unit for RPPA experiments; and Angie Fawkes and Lee Murphy at the Wellcome Trust Clinical Research Facility for RNA-sequencing. We thank our colleagues Mitchell Foster, Emily Webb, Virginia Alvarez Garcia, Jayne Culley, Niamh McGivern, Alfonso Bolado Carrancio and Roza Masalmeh for their valuable input and discussions surrounding our work. This work was funded by Cancer Research UK and the Medical Research Council. This work was supported by a joint Cancer Research UK (A28596) and The Brain Tumour Charity (GN-000676) funded Brain Tumour award to N.O.C and M.C.F. and a Cancer Research UK grant (A24837) awarded to V.B. and M.C.F, and a Cancer Research UK grant (A28592) awarded to S.M.P.

## Author Contributions

M.C.F. and V.B. co-ordinated the project; A.E.P.L., M.S.R., A.N.P., J.C.D., A.B., N.O.C., S.M.P., V.G.B. and M.C.F. designed the experiments and interpreted the results; A.E.P.L., M.S.R., A.N.P and M.T.M. performed the experiments; L.A., A.T.D., and R.L.M. produced and provided essential materials; M.C.F., V.B., S.M.P., N.O.C. and A.B. contributed to supervision; A.E.P.L. analysed the data and prepared figures; A.E.P.L. and M.F. wrote the paper. All authors read, commented on, and approved the manuscript.

---

**Supplementary Figure 1:**
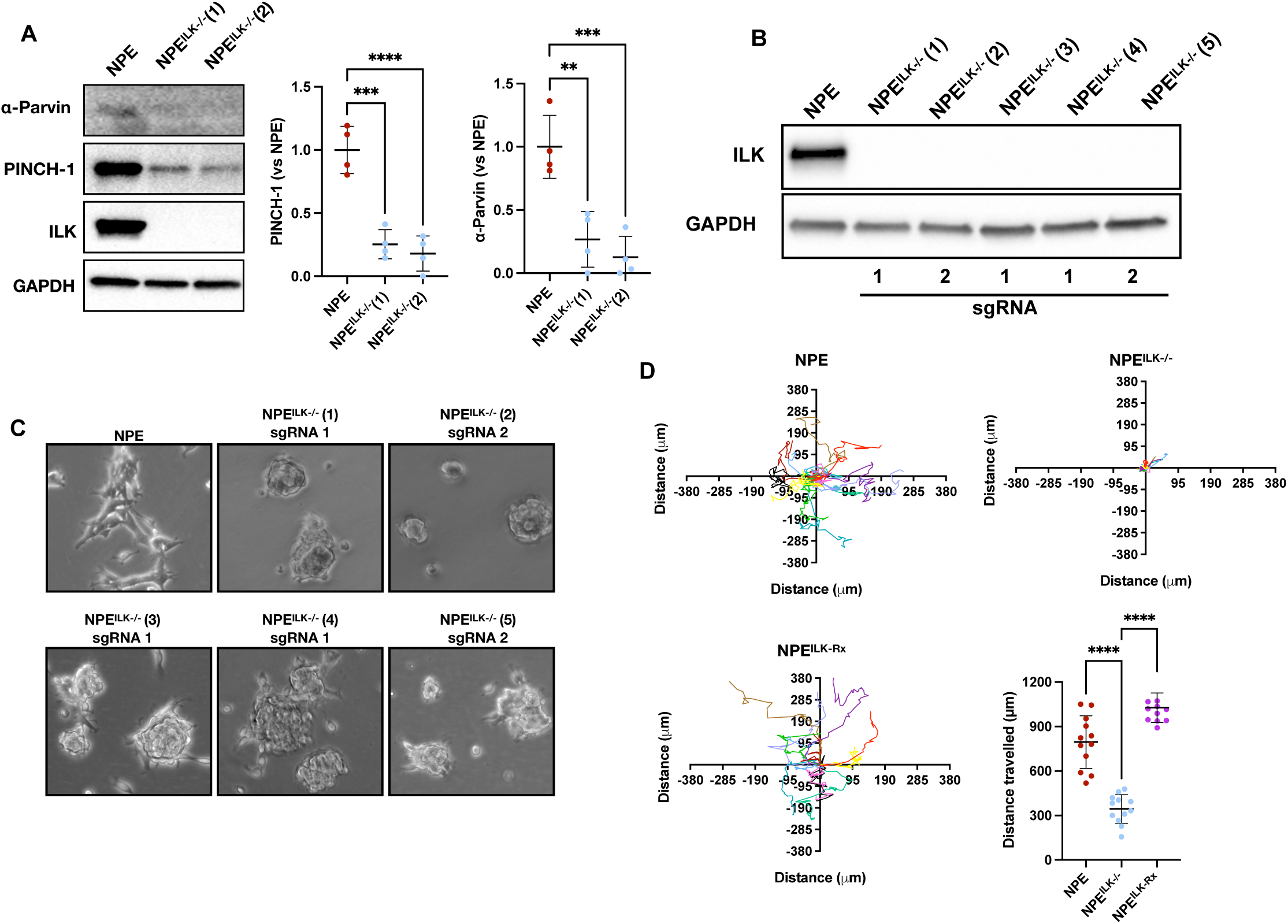
ILK controls morphology and is required for effective motility of transformed NSCs. **(A)** Western blotting for ILK and its binding partners PINCH1 and ɑ-Parvin in NPE cells and a pair of ILK-deficient NPE^ILK−/−^ clones (n = 4; \*\**P* < 0.01, \*\*\**P* < 0.005, \*\*\*\**P* < 0.001; One-way ANOVA with Dunnett’s multiple comparisons test). **(B)** Western blotting of NPE cells and 5 individual NPE^ILK−/−^ clones. sgRNA used to produce each clone is indicated. Clone 1 was used for ILK re-expression and further study. **(C)** Representative phase-contrast micrographs of NPE cells and 5 individual NPE^ILK−/−^ clones. sgRNA used to produce each clone is indicated. Clone 1 was used for ILK re-expression and further study. **(C)** Top left, top right and bottom left: Migration traces of randomly selected single NPE, NPE^ILK−/−^ and NPE^ILK-Rx^ cells. Each migration trace is centred on (0,0) (n = 12 representative single cells from 3 experiments per cell line). Bottom right: Quantification of total distance travelled by NPE, NPE^ILK−/−^ and NPE^ILK-Rx^ cells over 48 h (n = 12 representative single cells from 3 experiments; **** *P* < 0.001, one-way ANOVA with Tukey’s multiple comparisons test).

**Supplementary Figure 2:**
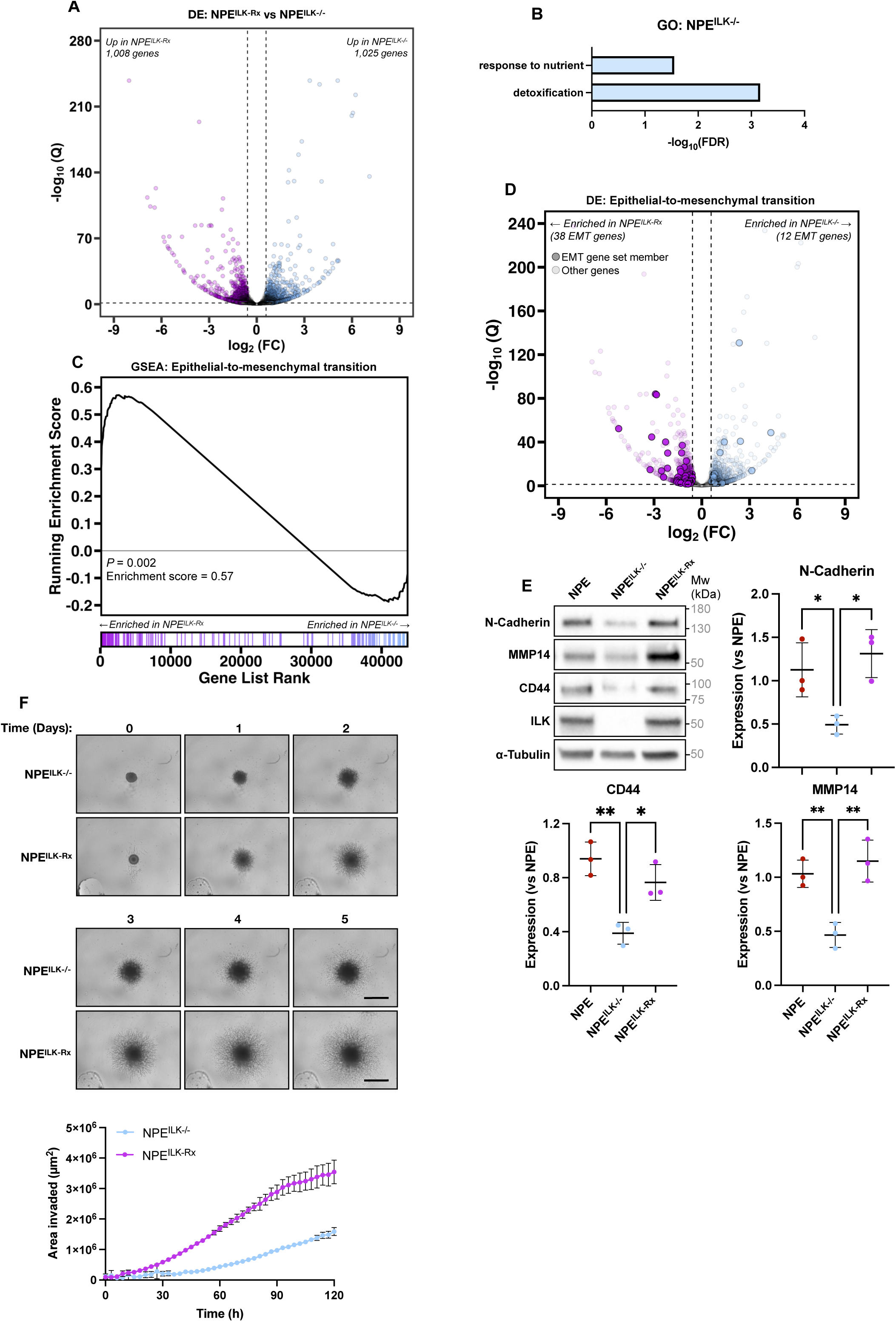
ILK is required for maintenance of mesenchymal state and stimulates GBM stem cell invasion. **(A)** Volcano plot of gene expression changes in NPE^ILK−/−^ vs NPE^ILK-Rx^ cells (n = 4; “DE” = differential expression) (Cutoffs shown of 1.5-fold expression change and false discovery rate (FDR)-adjusted *P* value < 0.05, Student’s *t*-test). **(B)** Gene ontology (GO) term overrepresentation analysis of statistically significantly upregulated genes (FDR < 0.05; fold change > 2) in NPE^ILK−/−^ *vs* NPE^ILK-Rx^ cells. **(C)** Gene set enrichment analysis (GSEA) performed on a 194-member epithelial-to-mesenchymal transition (EMT) gene set in NPE^ILK-Rx^ vs NPE^ILK−/−^ cells (n = 4). **(D)** Volcano plot of expression of 194 EMT-regulating genes in NPE^ILK-Rx^ vs NPE^ILK−/−^ cells. Shaded points indicate positions of EMT-regulating genes; unshaded points indicate positions of other genes (n = 4; “DE” = differential expression) (Cutoffs shown of 1.5-fold expression change and false discovery rate (FDR)-adjusted *P* value < 0.05, Student’s *t*-test). **(E)** Top left: Western blotting analysis of expression of indicated EMT-associated proteins in the indicated cell lines. Right and bottom: quantification of left (n = 3; ** *P* < 0.01, one-way ANOVA with Tukey’s multiple comparison test). **(F)** Top, middle: Representative phase-contrast micrographs of matrigel-embedded neurospheres derived from the indicated cell lines and at the indicated time-points (scale bar = 100 µm). Bottom left: quantification of invaded area (n = 3 independent experiments with 12 neurospheres per cell line each).

**Supplementary Figure 3:**
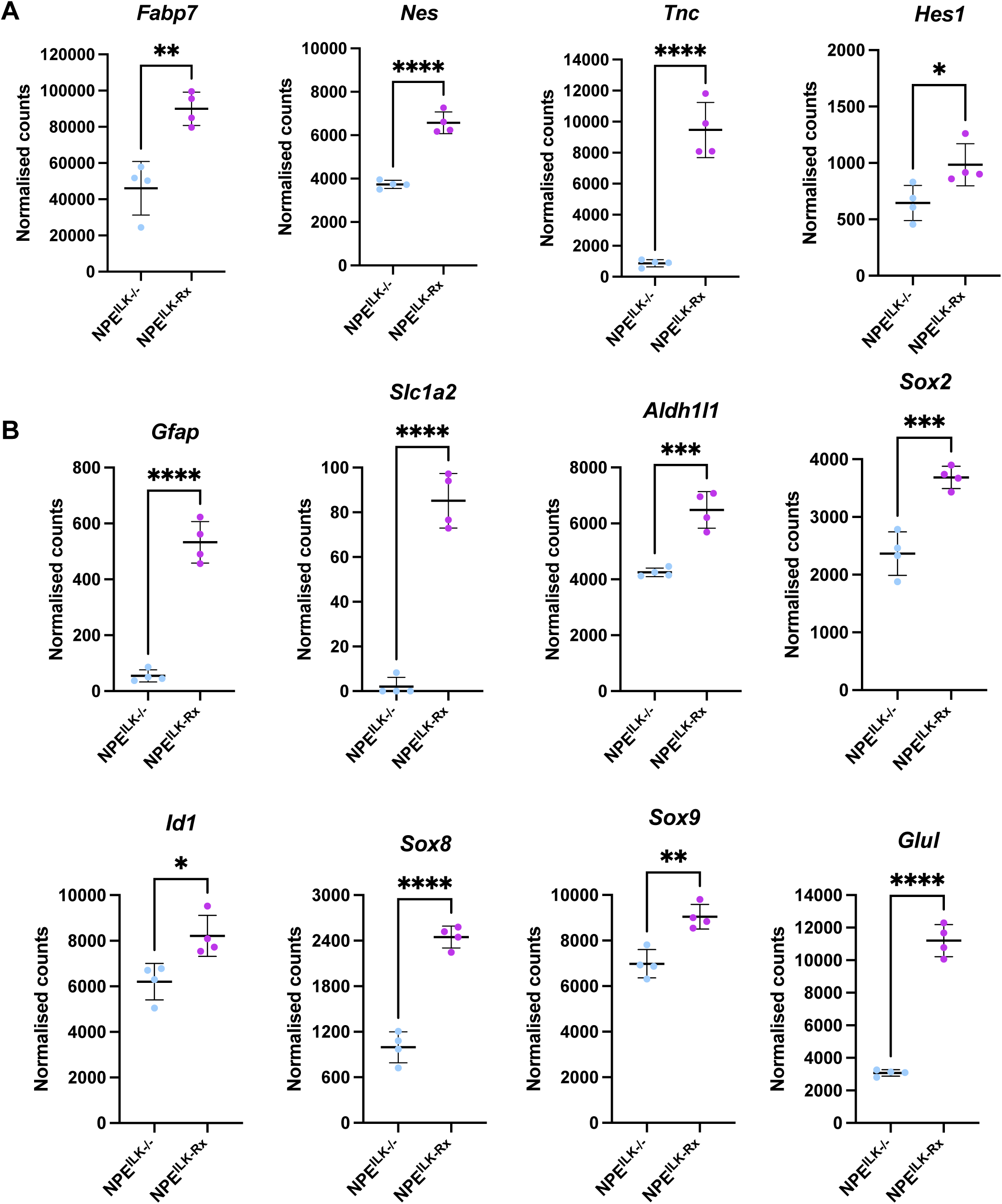
ILK depletion causes loss of consensus radial glia genes and inhibits induction of astrocyte-related genes following BMP-4 treatment. **(A)** RNA-seq expression of the indicated radial glia genes in NPE^ILK−/−^ and NPE^ILK-Rx^ cells (n = 4; \**P* < 0.05, \*\**P* < 0.01, \*\*\*\**P* < 0.001; Student’s two-tailed *t*-test). **(B)** RNA-seq expression of the indicated radial glia and astrocyte genes (*Gfap*, *Aldh1l1* and *Slc1a2*), transcriptional regulators involved in astrocyte differentiation (*Id1*, *Sox8* and *Sox9*), glutamine synthase (*Glul*), and *Sox2* in the indicated cell lines treated with BMP-4 for 24 h (n = 4; \**P* < 0.05, \*\**P* < 0.01, \*\*\**P* < 0.005, \*\*\*\**P* < 0.001; Student’s two-tailed *t*-test).

**Supplementary Figure 4:**
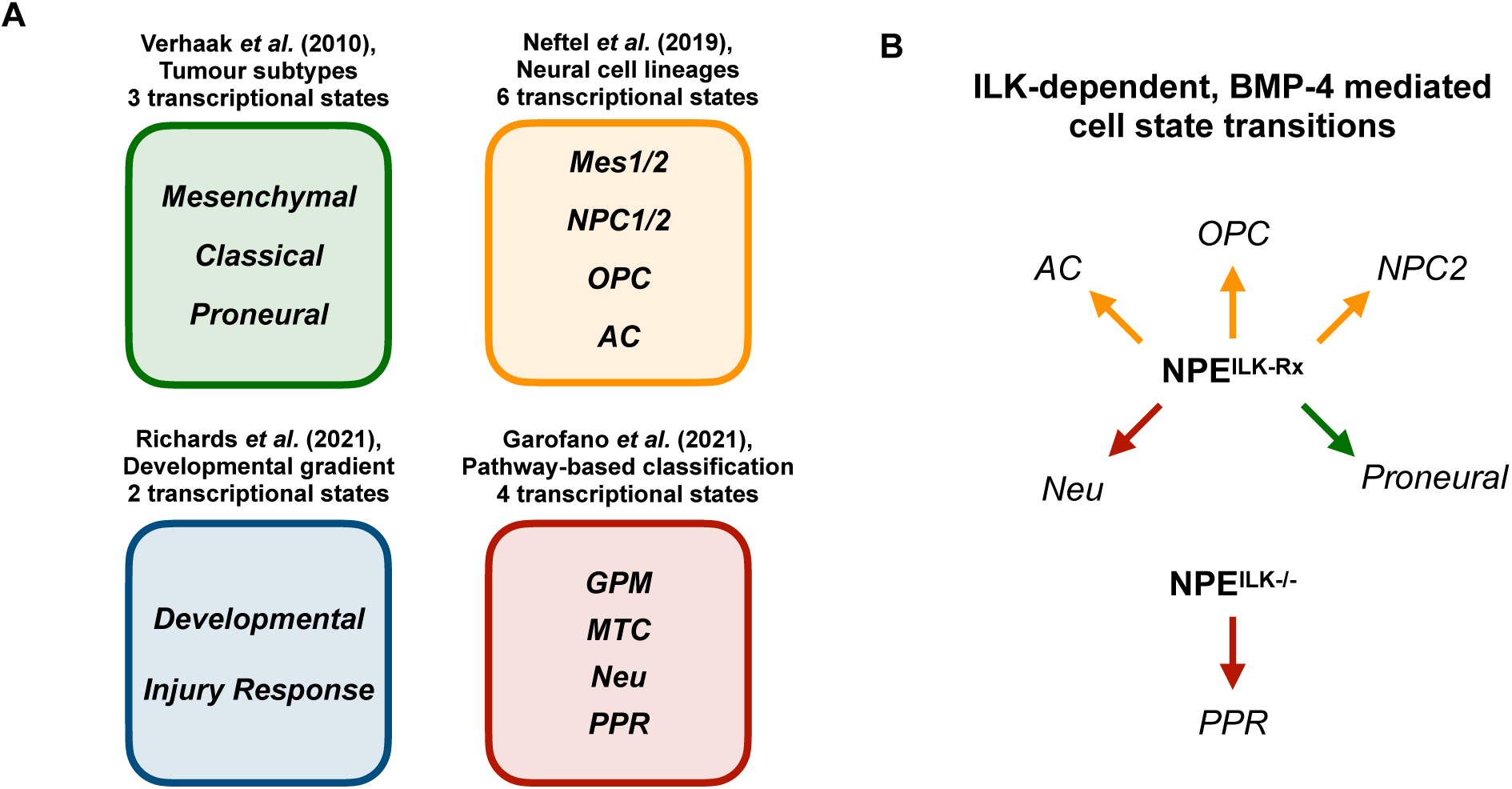
ILK expression facilitates diverse cell state transitions in BMP-4-treated GBM stem cells. **(A)** Graphical summary of literature-reported GBM cell states used in this study. **(B)** Schematic of ILK-mediated GBM cell state transitions in NPE^ILK-Rx^ and NPE^ILK−/−^ cells treated with BMP-4, related to Figure 2G. Arrow colour indicates the study that described each cell state.

**Supplementary Figure 5:**
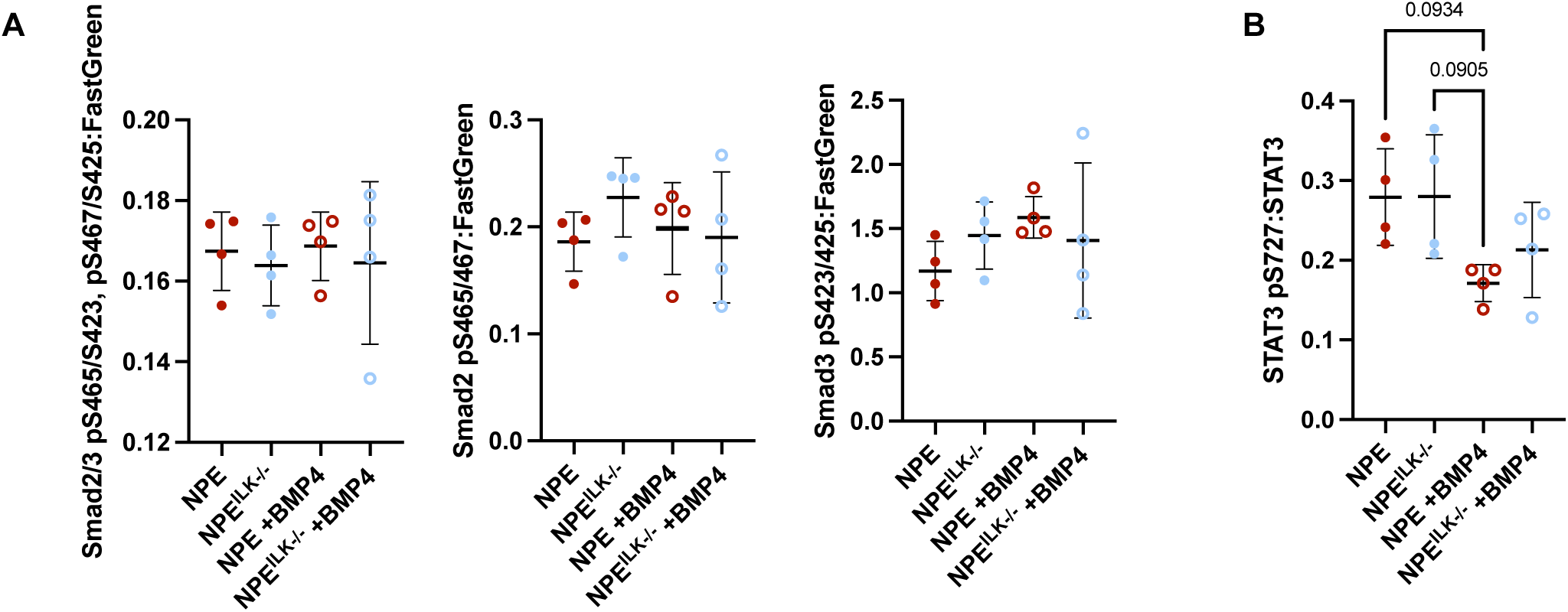
ILK does not affect Smad2/3 or STAT3 pS727 signalling in transformed NSCs. **(A)** Phosphorylation of the indicated Smad2/3 sites assayed by RPPA and normalised against the FastGreen total protein stain in NPE and NPE^ILK−/−^ cells either without treatment or following treatment with BMP-4 (n = 4; one-way ANOVA with Tukey’s multiple comparisons test). **(B)** Phosphorylation of STAT3 pS727, normalised against total STAT3 expression assayed by RPPA, in NPE and NPE^ILK−/−^ cells either untreated or treated with BMP-4 (n = 4; one-way ANOVA with Tukey’s multiple comparisons test).

**Supplementary Figure 6:**
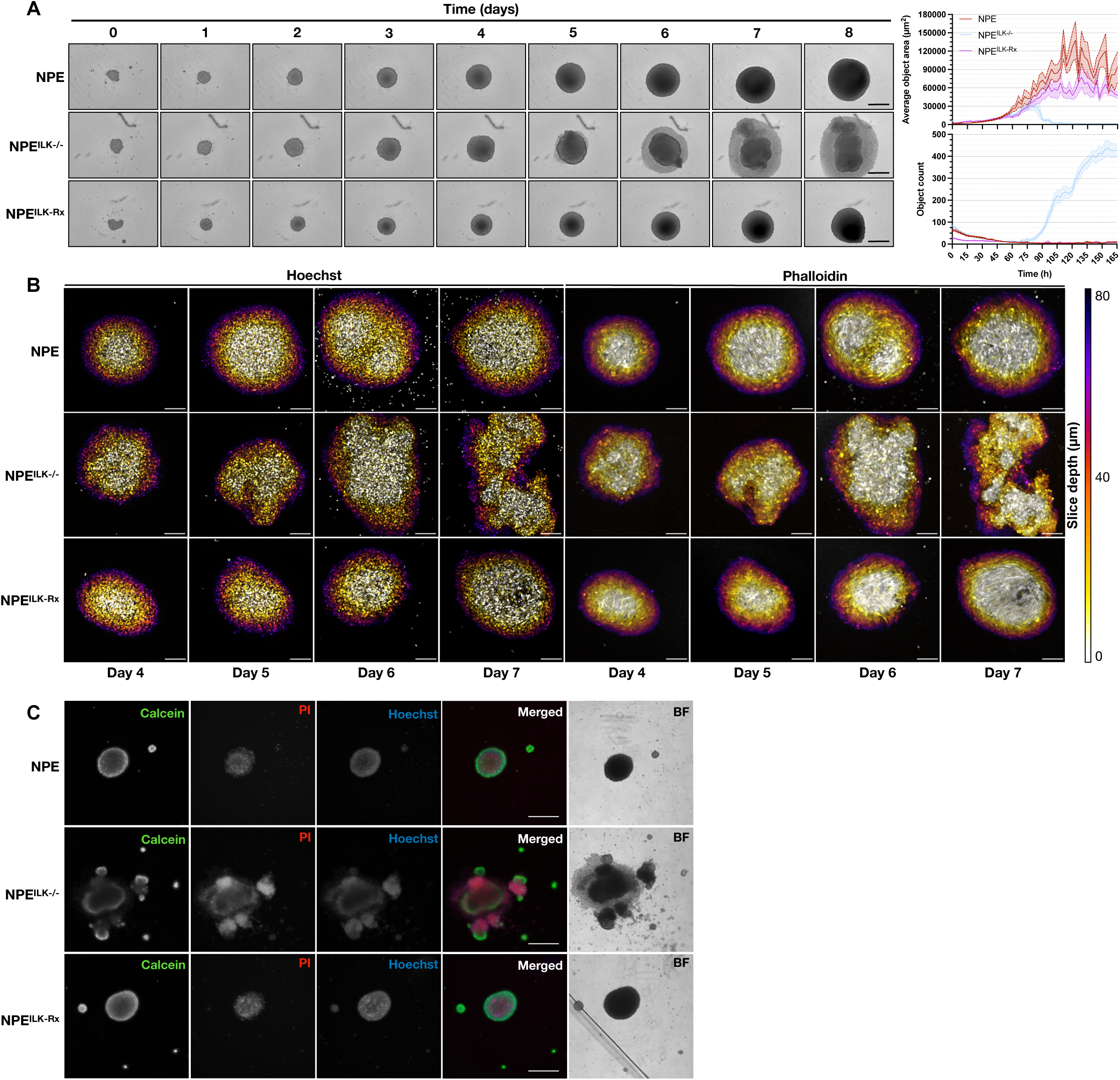
ILK regulates GBM neurosphere integrity. **(A)** Representative time-lapse images of NPE, NPE^ILK−/−^ and NPE^ILK-Rx^ neurospheres. Upper right: Quantification of average object area in images of neurospheres; lower right: quantification of total object count in images of neurospheres (n = 32 neurospheres per cell line). Scale bar = 500 µm. **(B)** Representative fluorescent images of NPE, NPE^ILK−/−^ and NPE^ILK-Rx^ neurospheres at the indicated time points staining nuclei (Hoechst-33342) and actin (phalloidin) (n = 12 neurospheres per cell line). Colours indicate Z-depth, with brighter colours being closer to the first imaging plane. Scale bar = Scale bar = 100 µm. **(C)** Representative fluorescent images of NPE, NPE^ILK−/−^ and NPE^ILK-Rx^ neurospheres at day 7 stained with calcein-AM (green; live cells), propidium iodide (red; PI; dead cells), or hoechst-33342 (blue; nuclei; all cells) (n = 8 neurospheres per cell line). Scale bar = 500 µm.

**Supplementary Figure 7:**
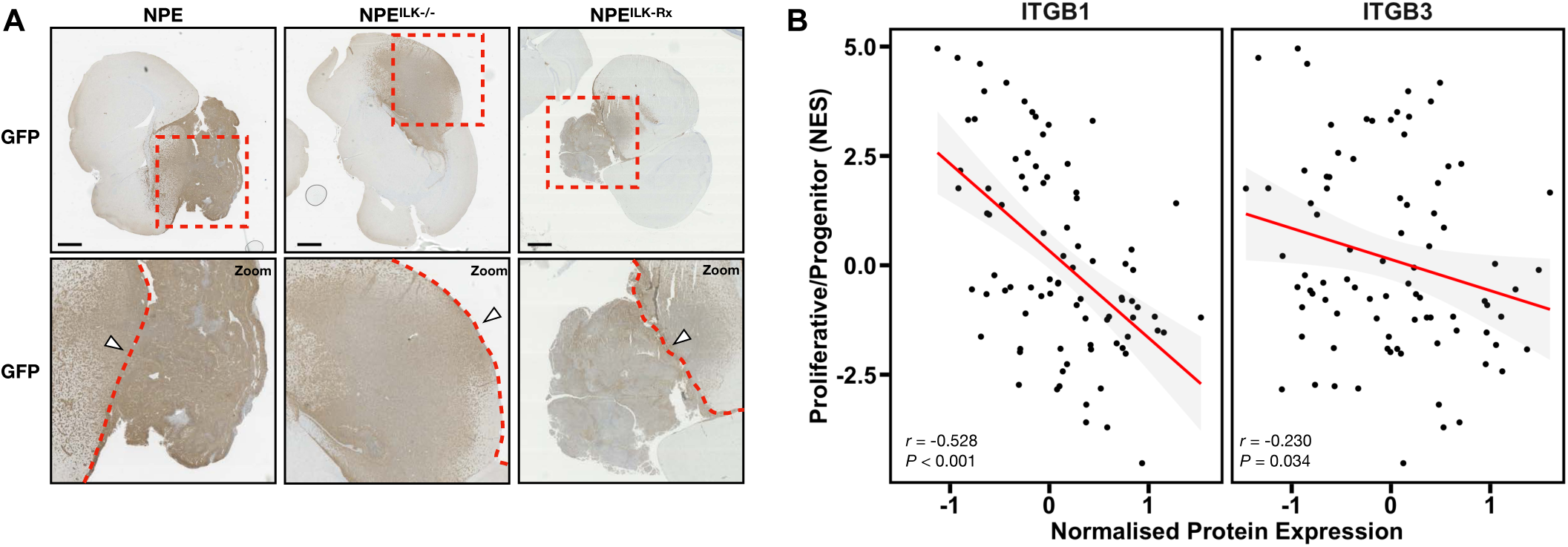
ILK affects tumour morphology and its integrin binding partners correlate with PPR signature expression. **(A)** Top panels: Representative immunohistochemistry images of GFP staining in brain slices from mice intracranially injected with 5,000 NPE^ILK−/−^ or NPE^ILK-Rx^ cells (images representative of 4 mice). Lower panels: magnified view of regions indicated by red boxes (top panels); dashed red lines mark approximate areas of brain periphery and white arrowheads mark approximate injection sites, with NPE and NPE^ILK-Rx^ tumours demonstrating outgrowth from injection sites. **(B)** Correlation between expression of β1 integrin (ITGB1) and β3 integrin (ITGB3) protein expression and PPR signature normalised enrichment scores (NES). Red trendline shown; shaded grey area indicates standard error (*r*: Pearson correlation coefficient).

## References

1. Davis, M.E. Glioblastoma: Overview of Disease and Treatment. Clin J Oncol Nurs 20, S2–8 (2016).

2. Robertson, F.L., Marques-Torrejon, M.A., Morrison, G.M. & Pollard, S.M. Experimental models and tools to tackle glioblastoma. Dis Model Mech 12 (2019).

3. Tamimi, A.F. & Juweid, M. Epidemiology and Outcome of Glioblastoma, in Glioblastoma. (ed. S. De Vleeschouwer) (Brisbane (AU); 2017).

4. Noroxe, D.S., Poulsen, H.S. & Lassen, U. Hallmarks of glioblastoma: a systematic review. ESMO Open 1, e000144 (2016).

5. Lan, X. et al. Fate mapping of human glioblastoma revea ls an invariant stem cell hierarchy. Nature 549, 227–232 (2017).

6. Richards, L.M. et al. Gradient of Developmental and Injury Response transcriptional states defines functional vulnerabilities underpinning glioblastoma heterogeneity. Nat Cancer 2, 157–173 (2021).

7. Qin, J. & Wu, C. ILK: a pseudokinase in the center stage of cell-matrix adhesion and signaling. Curr Opin Cell Biol 24, 607–613 (2012).

8. Paolillo, M. & Schinelli, S. Integrins and Exosomes, a Dangerous Liaison in Cancer Progression. Cancers (Basel) 9 (2017).

9. Desgrosellier, J.S. & Cheresh, D.A. Integrins in cancer: biological implications and therapeutic opportunities. Nat Rev Cancer 10, 9–22 (2010).

10. Yu, Q. et al. Fibronectin Promotes the Malignancy of Glioma Stem-Like Cells Via Modulation of Cell Adhesion, Differentiation, Proliferation and Chemoresistance. Front Mol Neurosci 11, 130 (2018).

11. Lathia, J.D. et al. Laminin alpha 2 enables glioblastoma stem cell growth. Ann Neurol 72, 766–778 (2012).

12. Durbeej, M. Laminins. Cell Tissue Res 339, 259–268 (2010).

13. Rezk, R. et al. Spatial heterogeneity of cell-matrix adhesive forces predicts human glioblastoma migration. Neurooncol Adv 2, vdaa081 (2020).

14. Horton, E.R. et al. Definition of a consensus integrin adhesome and its dynamics during adhesion complex assembly and disassembly. Nat Cell Biol 17, 1577–1587 (2015).

15. Gangoso, E. et al. Glioblastomas acquire myeloid-affiliated transcriptional programs via epigenetic immunoediting to elicit immune evasion. Cell 184, 2454–2470 e2426 (2021).

16. Cheng, L. et al. Elevated invasive potential of glioblastoma stem cells. Biochem Biophys Res Commun 406, 643–648 (2011).

17. Wenger, A., Larsson, S. & Caren, H. Patient-derived cells modeling pediatric glioma. Aging (Albany NY) 9, 1353–1354 (2017).

18. Venere, M., Fine, H.A., Dirks, P.B. & Rich, J.N. Cancer stem cells in gliomas: identifying and understanding the apex cell in cancer’s hierarchy. Glia 59, 1148–1154 (2011).

19. Lathia, J.D., Mack, S.C., Mulkearns-Hubert, E.E., Valentim, C.L. & Rich, J.N. Cancer stem cells in glioblastoma. Genes Dev 29, 1203–1217 (2015).

20. Lathia, J.D. et al. Direct in vivo evidence for tumor propagation by glioblastoma cancer stem cells. PLoS One 6, e24807 (2011).

21. Caren, H. et al. Glioblastoma Stem Cells Respond to Differentiation Cues but Fail to Undergo Commitment and Terminal Cell-Cycle Arrest. Stem Cell Reports 5, 829–842 (2015).

22. Pollard, S.M. In vitro expansion of fetal neural progenitors as adherent cell lines. Methods Mol Biol 1059, 13–24 (2013).

23. Pollard, S.M., Conti, L., Sun, Y., Goffredo, D. & Smith, A. Adherent neural stem (NS) cells from fetal and adult forebrain. Cereb Cortex 16 Suppl 1, i112–120 (2006).

24. Gimple, R.C., Bhargava, S., Dixit, D. & Rich, J.N. Glioblastoma stem cells: lessons from the tumor hierarchy in a lethal cancer. Genes Dev 33, 591–609 (2019).

25. Stanchi, F. et al. Molecular dissection of the ILK-PINCH-parvin triad reveals a fundamental role for the ILK kinase domain in the late stages of focal-adhesion maturation. J Cell Sci 122, 1800–1811 (2009).

26. Chiswell, B.P. et al. Structural basis of competition between PINCH1 and PINCH2 for binding to the ankyrin repeat domain of integrin-linked kinase. J Struct Biol 170, 157–163 (2010).

27. McBeath, R., Pirone, D.M., Nelson, C.M., Bhadriraju, K. & Chen, C.S. Cell shape, cytoskeletal tension, and RhoA regulate stem cell lineage commitment. Dev Cell 6, 483–495 (2004).

28. Barker, C.G. et al. Identification of phenotype-specific networks from paired gene expression-cell shape imaging data. Genome Res 32, 750–765 (2022).

29. Widmaier, M., Rognoni, E., Radovanac, K., Azimifar, S.B. & Fassler, R. Integrin-linked kinase at a glance. J Cell Sci 125, 1839–1843 (2012).

30. Liberzon, A. et al. The Molecular Signatures Database (MSigDB) hallmark gene set collection. Cell Syst 1, 417–425 (2015).

31. Zeisberg, M. & Neilson, E.G. Biomarkers for epithelial-mesenchymal transitions. J Clin Invest 119, 1429–1437 (2009).

32. Xu, H. et al. The role of CD44 in epithelial-mesenchymal transition and cancer development. Onco Targets Ther 8, 3783–3792 (2015).

33. Carro, M.S. et al. The transcriptional network for mesenchymal transformation of brain tumours. Nature 463, 318–325 (2010).

34. Liddelow, S.A. et al. Neurotoxic reactive astrocytes are induced by activated microglia. Nature 541, 481–487 (2017).

35. Cahoy, J.D. et al. A transcriptome database for astrocytes, neurons, and oligodendrocytes: a new resource for understanding brain development and function. J Neurosci 28, 264–278 (2008).

36. Zhong, S. et al. A single-cell RNA-seq survey of the developmental landscape of the human prefrontal cortex. Nature 555, 524–528 (2018).

37. Nowakowski, T.J. et al. Spatiotemporal gene expression trajectories reveal developmental hierarchies of the human cortex. Science 358, 1318–1323 (2017).

38. Pollen, A.A. et al. Molecular identity of human outer radial glia during cortical development. Cell 163, 55–67 (2015).

39. Anthony, T.E., Klein, C., Fishell, G. & Heintz, N. Radial glia serve as neuronal progenitors in all regions of the central nervous system. Neuron 41, 881–890 (2004).

40. Eze, U.C., Bhaduri, A., Haeussler, M., Nowakowski, T.J. & Kriegstein, A.R. Single-cell atlas of early human brain development highlights heterogeneity of human neuroepithelial cells and early radial glia. Nat Neurosci 24, 584–594 (2021).

41. Tabata, H. Diverse subtypes of astrocytes and their development during corticogenesis. Front Neurosci 9, 114 (2015).

42. Chandrasekaran, A., Avci, H.X., Leist, M., Kobolak, J. & Dinnyes, A. Astrocyte Differentiation of Human Pluripotent Stem Cells: New Tools for Neurological Disorder Research. Front Cell Neurosci 10, 215 (2016).

43. Neftel, C. et al. An Integrative Model of Cellular States, Plasticity, and Genetics for Glioblastoma. Cell 178, 835–849 e821 (2019).

44. Patel, A.P. et al. Single-cell RNA-seq highlights intratumoral heterogeneity in primary glioblastoma. Science 344, 1396–1401 (2014).

45. Eng, L.F. Glial fibrillary acidic protein (GFAP): the major protein of glial intermediate filaments in differentiated astrocytes. J Neuroimmunol 8, 203–214 (1985).

46. Guttenplan, K.A. & Liddelow, S.A. Astrocytes and microglia: Models and tools. J Exp Med 216, 71–83 (2019).

47. Johnson, K. et al. Gfap-positive radial glial cells are an essential progenitor population for later-born neurons and glia in the zebrafish spinal cord. Glia 64, 1170–1189 (2016).

48. Metz, E.P. et al. Tumor quiescence: elevating SOX2 in diverse tumor cell types downregulates a broad spectrum of the cell cycle machinery and inhibits tumor growth. BMC Cancer 20, 941 (2020).

49. Wang, Y. et al. SOX2 is essential for astrocyte maturation and its deletion leads to hyperactive behavior in mice. Cell Rep 41, 111842 (2022).

50. Sun, W. et al. SOX9 Is an Astrocyte-Specific Nuclear Marker in the Adult Brain Outside the Neurogenic Regions. J Neurosci 37, 4493–4507 (2017).

51. Takouda, J., Katada, S., Imamura, T., Sanosaka, T. & Nakashima, K. SoxE group transcription factor Sox8 promotes astrocytic differentiation of neural stem/precursor cells downstream of Nfia. Pharmacol Res Perspect 9, e00749 (2021).

52. Bjornsen, L.P., Hadera, M.G., Zhou, Y., Danbolt, N.C. & Sonnewald, U. The GLT-1 (EAAT2; slc1a2) glutamate transporter is essential for glutamate homeostasis in the neocortex of the mouse. J Neurochem 128, 641–649 (2014).

53. Beyer, F., Ludje, W., Karpf, J., Saher, G. & Beckervordersandforth, R. Distribution of Aldh1L1-CreER(T2) Recombination in Astrocytes Versus Neural Stem Cells in the Neurogenic Niches of the Adult Mouse Brain. Front Neurosci 15, 713077 (2021).

54. Nam, H.S. & Benezra, R. High levels of Id1 expression define B1 type adult neural stem cells. Cell Stem Cell 5, 515–526 (2009).

55. Hanzelmann, S., Castelo, R. & Guinney, J. GSVA: gene set variation analysis for microarray and RNA-seq data. BMC Bioinformatics 14, 7 (2013).

56. Garofano, L. et al. Pathway-based classification of glioblastoma uncovers a mitochondrial subtype with therapeutic vulnerabilities. Nat Cancer 2, 141–156 (2021).

57. Verhaak, R.G. et al. Integrated genomic analysis identifies clinically relevant subtypes of glioblastoma characterized by abnormalities in PDGFRA, IDH1, EGFR, and NF1. Cancer Cell 17, 98–110 (2010).

58. Migliozzi, S. et al. Integrative multi-omics networks identify PKCdelta and DNA-PK as master kinases of glioblastoma subtypes and guide targeted cancer therapy. Nat Cancer 4, 181–202 (2023).

59. Mort, R.L. et al. Fucci2a: a bicistronic cell cycle reporter that allows Cre mediated tissue specific expression in mice. Cell Cycle 13, 2681–2696 (2014).

60. Wagner, T.U. Bone morphogenetic protein signaling in stem cells--one signal, many consequences. FEBS J 274, 2968–2976 (2007).

61. Osnato, A. et al. TGFbeta signalling is required to maintain pluripotency of human naive pluripotent stem cells. Elife 10 (2021).

62. Yang, J. & Jiang, W. The Role of SMAD2/3 in Human Embryonic Stem Cells. Front Cell Dev Biol 8, 653 (2020).

63. Rajan, P., Panchision, D.M., Newell, L.F. & McKay, R.D. BMPs signal alternately through a SMAD or FRAP-STAT pathway to regulate fate choice in CNS stem cells. J Cell Biol 161, 911–921 (2003).

64. Bonni, A. et al. Regulation of gliogenesis in the central nervous system by the JAK-STAT signaling pathway. Science 278, 477–483 (1997).

65. Hong, S. & Song, M.R. STAT3 but not STAT1 is required for astrocyte differentiation. PLoS One 9, e86851 (2014).

66. Huang, G., Yan, H., Ye, S., Tong, C. & Ying, Q.L. STAT3 phosphorylation at tyrosine 705 and serine 727 differentially regulates mouse ESC fates. Stem Cells 32, 1149–1160 (2014).

67. Mogul, A., Corsi, K. & McAuliffe, L. Baricitinib: The Second FDA-Approved JAK Inhibitor for the Treatment of Rheumatoid Arthritis. Ann Pharmacother 53, 947–953 (2019).

68. Hubbard, J.A., Hsu, M.S., Seldin, M.M. & Binder, D.K. Expression of the Astrocyte Water Channel Aquaporin-4 in the Mouse Brain. ASN Neuro 7 (2015).

69. Wang, L.B. et al. Proteogenomic and metabolomic characterization of human glioblastoma. Cancer Cell 39, 509–528 e520 (2021).

70. Vaynberg, J. et al. Non-catalytic signaling by pseudokinase ILK for regulating cell adhesion. Nat Commun 9, 4465 (2018).

71. Porcheri, C., Suter, U. & Jessberger, S. Dissecting integrin-dependent regulation of neural stem cell proliferation in the adult brain. J Neurosci 34, 5222–5232 (2014).

72. Garcia-Diaz, C. et al. Glioblastoma cell fate is differentially regulated by the microenvironments of the tumor bulk and infiltrative margin. Cell Rep 42, 112472 (2023).

73. Jain, A. et al. Guiding intracortical brain tumour cells to an extracortical cytotoxic hydrogel using aligned polymeric nanofibres. Nat Mater 13, 308–316 (2014).

74. Lee, S.Y. Temozolomide resistance in glioblastoma multiforme. Genes Dis 3, 198–210 (2016).

75. Schneider, C.A., Rasband, W.S. & Eliceiri, K.W. NIH Image to ImageJ: 25 years of image analysis. Nat Methods 9, 671–675 (2012).

76. Slaymaker, I.M. et al. Rationally engineered Cas9 nucleases with improved specificity. Science 351, 84–88 (2016).

77. Rizzo, M.A., Springer, G.H., Granada, B. & Piston, D.W. An improved cyan fluorescent protein variant useful for FRET. Nat Biotechnol 22, 445–449 (2004).

78. Kanda, T., Sullivan, K.F. & Wahl, G.M. Histone-GFP fusion protein enables sensitive analysis of chromosome dynamics in living mammalian cells. Curr Biol 8, 377–385 (1998).

79. Avalle, L. et al. Liver-Specific siRNA-Mediated Stat3 or C3 Knockdown Improves the Outcome of Experimental Autoimmune Myocarditis. Mol Ther Methods Clin Dev 18, 62–72 (2020).

80. Bankhead, P. et al. QuPath: Open source software for digital pathology image analysis. Sci Rep 7, 16878 (2017).

81. Macleod, K.G., Serrels, B. & Carragher, N.O. Reverse Phase Protein Arrays and Drug Discovery. Methods Mol Biol 1647, 153–169 (2017).

82. Love, M.I., Huber, W. & Anders, S. Moderated estimation of fold change and dispersion for RNA-seq data with DESeq2. Genome Biol 15, 550 (2014).

83. Wang, J., Vasaikar, S., Shi, Z., Greer, M. & Zhang, B. WebGestalt 2017: a more comprehensive, powerful, flexible and interactive gene set enrichment analysis toolkit. Nucleic Acids Res 45, W130–W137 (2017).

84. Subramanian, A. et al. Gene set enrichment analysis: a knowledge-based approach for interpreting genome-wide expression profiles. Proc Natl Acad Sci U S A 102, 15545–15550 (2005).

85. Wickham, H., Edn. Second edition (Springer, Switzerland; 2016).

86. Gu, Z., Eils, R. & Schlesner, M. Complex heatmaps reveal patterns and correlations in multidimensional genomic data. Bioinformatics 32, 2847–2849 (2016).

